# Targeting AAV vectors to the CNS via *de novo* engineered capsid-receptor interactions

**DOI:** 10.1101/2022.10.31.514553

**Authors:** Qin Huang, Albert T. Chen, Ken Y. Chan, Hikari Sorensen, Andrew J. Barry, Bahar Azari, Thomas Beddow, Qingxia Zheng, Binhui Zhao, Isabelle G. Tobey, Fatma-Elzahraa Eid, Yujia A. Chan, Benjamin E. Deverman

## Abstract

Viruses have evolved the ability to bind and enter cells through interactions with a wide variety of host cell macromolecules. Here, we screened for AAV capsids that bind two host cell proteins expressed on the mouse blood-brain barrier, LY6A or the related protein LY6C1. Introducing interactions with either protein target generated hundreds of capsids with dramatically enhanced central nervous system (CNS) tropisms. In contrast to the AAV-PHP.B capsid family, which interacts with LY6A and only exhibits its enhanced CNS tropism in a subset of mouse strains, the capsids that engage LY6C1 maintain their CNS tropism in BALB/cJ mice. Compared to conventional *in vivo* screens for CNS cell transducing capsids, a single round of protein target binding screening recovered significantly more capsids with enhanced performance that were validated in subsequent *in vivo* screens. Moreover, the initial screening round generated reproducible and quantitative target binding data that enabled the efficient machine learning-guided generation of more diverse target-specific capsids. This work demonstrates that AAV capsids can be directly targeted to specific proteins to generate potent gene delivery vectors with known mechanisms of action and predictable tropisms.

## Introduction

Gene therapy with recombinant adeno-associated viruses (AAVs) shows promise for treating diseases at their root genetic cause but remains constrained by the inefficiency of delivery to disease-relevant organs and cell types. Natural AAV capsids can be modified to produce vectors with dramatically improved *in vivo* tropisms. An effective engineering strategy has been to generate diverse libraries of capsid variants via peptide insertions, and to subject these libraries to multiple rounds of *in vivo* selection to identify capsids with the desired properties such as CNS-wide transduction (Chan *et al*., 2017; Deverman *et al*., 2016; Nonnenmacher *et al*.,2021), brain vascular endothelium targeting (Körbelin *et al*., 2016; Krolak *et al*., 2022), retrograde transduction in the CNS (Tervo *et al*., 2016), transduction of human hepatocytes in a xenograft system (Lisowski *et al*.,2013), photoreceptor transduction (Dalkara *et al*., 2013), and muscle transduction (Tabebordbar *et al*., 2021; Weinmann *et al*., 2020). However, these screening efforts have been limited to function-focused approaches, where capsids are selected for a particular biodistribution or cell type tropism without discriminating for mechanism of action. The mechanism underlying the selected function must typically be elucidated via detailed downstream studies. As a result, the rare high-performance capsids identified in these screens may rely on mechanisms of action not conserved across host species (Hordeaux *et al*., 2018; Huang *et al*., 2019; Körbelin *et al*., 2016; Lisowski *et al*., 2013). Human cell or organoid models of increasing sophistication may provide new opportunities for human-relevant capsid engineering (Brown *et al*., 2019; Cho *et al*., 2017; Depla *et al*.,2020; Garita-Hernandez *et al*., 2020; Liang and Yoon, 2021; Sherman and Rossi, 2019). However, without a clear and preserved underlying mechanism of action, capsids selected *in vitro* may not retain their selected function *in vivo*.

Several research groups have attempted to circumvent this shortcoming by innovating mechanism-focused approaches, e.g., grafting independently characterized or engineered peptides (Shi and Bartlett, 2003; White *et al*., 2008; Yu *et al*., 2009) or proteins, such as DARPins or antibody fragments, onto the AAV capsid (Eichhoff *et al*., 2019; Hamann *et al*., 2021; Muik *et al*., 2017; Münch *et al*., 2015, 2013; Ponnazhagan *et al*., 2002; Reul *et al*., 2019). However, these grafting approaches do not select for optimal affinity in the context of the functional vector and may increase the complexity of manufacturing. Therefore, the majority of AAV capsid engineering efforts to date have continued to focus on *in vivo* selections.

In 2019, we and others reported that AAV-PHP.B (Deverman *et al*., 2016) and related capsids (Chan *et al*., 2017) could utilize a novel blood-brain barrier (BBB)-crossing mechanism by interacting with the LY6A protein on the surface of the brain endothelium of a subset of mouse strains (Hordeaux et al., 2019; Huang *et al*., 2019). Based on this finding, we were encouraged to develop a novel mechanism-focused approach that screens an AAV capsid library for variants that bind host proteins that are likely to translate into a desired *in vivo* tropism – in this case, BBB-crossing activity. As a proof-of-concept, we targeted two mouse CNS endothelium proteins, LY6A and LY6C1, and used pull-down assays to screen for AAVs capable of directly binding these target proteins *in vitro*. A large fraction of the capsids engineered to interact with LY6A or LY6C1 *in vitro* exhibited *in vivo* BBB-crossing activity that was enhanced relative to AAV9 and comparable to other reported capsids with improved CNS tropisms. Due to the conserved expression of LY6C1 across the CNS endothelium of commonly used mouse strains, capsids that engage LY6C1 have an enhanced CNS tropism in strains that are nonpermissive to AAV-PHP.B. Furthermore, this approach generated highly quantitative and reproducible data from a single round of screening, which enabled rapid motif identification and the generation of a diverse set of additional sequences via saturation mutagenesis and the use of a supervised variational auto-encoder (SVAE) ML model. Many of these additional variants were found to exhibit high levels of *in vivo* CNS transduction. This work demonstrates that AAV capsids can be systematically targeted to defined cell surface proteins to facilitate enhanced and predictable *in vivo* tropisms.

## Results

### A high-throughput purified protein assay identifies capsids selective for LY6A or LY6C1

To assess the potential of a mechanism-focused approach to develop capsids with enhanced CNS tropisms, we targeted two surface proteins present on brain vascular endothelial cells: LY6A, the known receptor for the AAV-PHP.B family of capsids, as a positive control, and a related protein, LY6C1, a novel target likewise highly expressed on CNS endothelial cells (Huang *et al*., 2019; Zhang *et al*.,2014). LY6C1 was selected based on the hypothesis that it may share LY6A’s ability to mediate AAV transport into the CNS, given that the LY6 family possesses a conserved protein structure and subcellular localization (Loughner *et al*., 2016). We generated LY6A and LY6C1 proteins as Fc fusions and used a magnetic bead-based pull-down assay to perform initial (Round 1) screens of two independently generated 7-mer-modified AAV9 libraries (random 7-mer amino acid sequences were inserted between residues 588–589 in VP1) - named Library 1 and Library 2, respectively - for variants that bind to LY6A-Fc, LY6C1-Fc, or an Fc-only control (Library 1 data is shown in Figure 1B–E; Library 2 data is shown in Figure 1—figure supplement 1). In the libraries, each capsid variant packaged its own capsid-encoding genome, allowing for the assessment of binding to the target by short-read, next-generation sequencing (NGS). The pull-down assays yielded reproducible binding scores for capsid variants across a wide dynamic range, with a high correlation of read depth-normalized counts between replicates (Figure 1B, Figure 1—figure supplement 2A–H).

**Figure 1.**
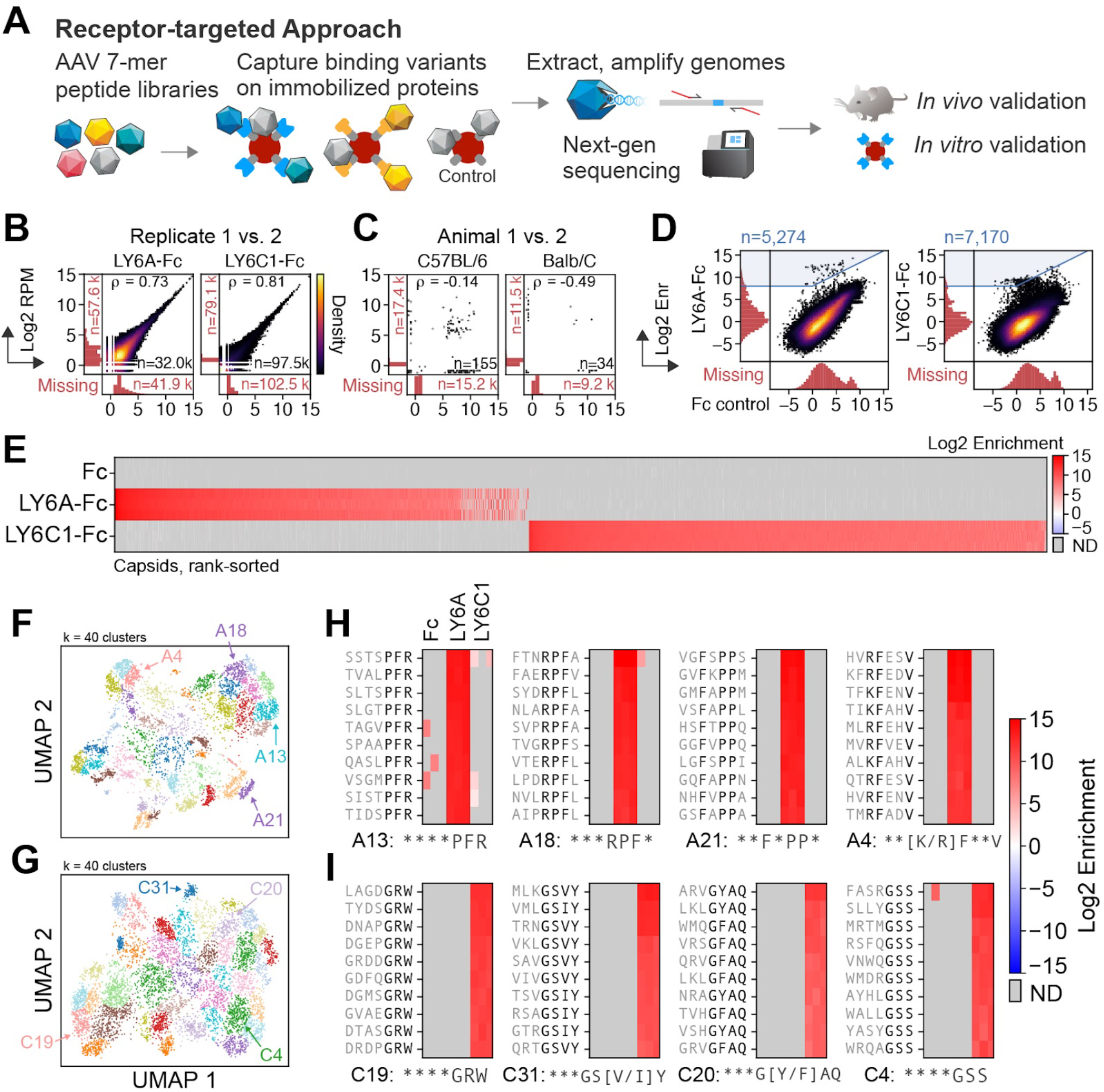
*In vitro* pull-down assays yield capsids that selectively bind LY6A or LY6C1. (**A**) A capsid library is screened for the ability to bind immobilized target Fc-fusion proteins. Bound capsid sequences are extracted and subjected to NGS. Hits are incorporated into a focused library for *in vivo* and *in vitro* validation. (**B,C**) The Pearson correlation of the log_2_ normalized read count (reads per million, RPM) are shown between biological replicates (n = 3, only one pair shown) (**B**) and between animals (n = 2) (**C**). Variants detected in one replicate or animal and not the other are shown in the marginal histograms. (**D**) The variant log_2_ enrichment (average RPM between replicates, normalized to the starting library RPM) plotted between the target (y-axis) and Fc-only control (x-axis) shows a majority of variants with nonspecific binding and a minority (blue highlighted region) with target-specific binding. The variants detected in one assay and not the other are shown in the marginal histograms. (**E**) The log_2_ enrichment of the selected variants highlighted in blue in (**D**) with each replicate’s enrichment plotted in separate rows (n = 3). ND = not detected. (**F,G**) The sequences that bound LY6A (**F**) or LY6C1 (**G**) from Libraries 1 and 2 were one-hot encoded, jointly projected with UMAP, and jointly clustered with a Gaussian mixture model (k = 40). **(H,I)**Four clusters for each target from (**F,G**) were manually selected based on whether there was a clear motif 2–4 amino acids in length that matched either existing reference sequence (LY6A-binding: ***PFR, ***RPF, LY6C1-binding: ***G[Y/F]AQ) or represented a motif not yet seen in published studies. Consensus motifs are defined per-position, with flexible amino acid residues (asterisks) and fixed residues (present in more than 40% of the cluster’s sequences; black letters).

To compare the pull-down assays to a conventional *in vivo* selection, we screened Library 2 for capsids that transduced the C57BL/6J and BALB/cJ CNS (n = 2 mice per strain) using transcribed capsid sequences as a functional readout (Krolak *et al*., 2022; Nonnenmacher *et al*., 2021). The vast majority of variants were detected in only one animal (Figure 1C, Figure 1—figure supplement 2I,J) as has been observed in other *in vivo* selection experiments (Nonnenmacher *et al*., 2021; Tabebordbar *et al*., 2021). In contrast, the pull-down assays yielded thousands of unique capsids that selectively bound the intended target but not the Fc-only control (Figure 1D, Figure 1—figure supplement 1A) or the other target, i.e., capsids selected for LY6C1 binding were not highly enriched for LY6A binding, and vice versa (Figure 1E).

### A wide array of sequence motifs are enriched by the pull-down assays

To assess the diversity of sequences enriched among LY6A- and LY6C1-binding 7-mers, we projected the protein-specific sequences highlighted in Figure 1D using UMAP (McInnes *et al*., 2018) and jointly clustered sequences from Library 1 and Library 2 with a Gaussian mixture model (k = 40) (Figure 1F,G; Figure 1—figure supplement 3A,B; Supplementary files 1–6). All clusters for both LY6A and LY6C1 have representatives from both libraries (Figure 1—figure supplement 3C,D). This consistency across independent libraries demonstrates that the approach can reproducibly detect thousands of unique capsid sequences with common sequence motifs. Inspection of the clusters of LY6A- and LY6C1-binding 7-mers revealed clear sequence motifs, typically 2–4 amino acids in length. Of these motifs, some were similar to previously published capsid sequences with CNS tropisms (Figure 1F–I, all clusters are shown in Figure 1—figure supplement 4 and Supplementary files 1–6). For example, clusters containing sequences similar to those of known LY6A-binding capsids were observed: AAV-PHP.B (TLAVPFK), clusters A5, 13, 33; AAV-PHP.B2 (SVSKPFL), clusters A14, 18, 32; and AAV-PHP.B3 (FTLTTPK), clusters A17, 35 (Supplementary file 7).

### *In vitro* selection for LY6A and LY6C1 binding yields capsids with enhanced tropisms predicted by target expression

To test whether the capsids selected to target LY6A or LY6C1 *in vitro* enable efficient BBB crossing, we generated a Round 2 library containing top hits from the initial Round 1 screen for LY6A and LY6C1 binding (n = 6.4K and 12.6K unique 7-mers, respectively). To compare the pull-down assay approach to conventional *in vivo* selections, we included all unique 7-mer sequences recovered following the Round 1 screening for expression in the CNS of C57BL/6J or BALB/cJ mice (n = 5.8K) (Figure 2A; only Library 2 was used in the Round 1 *in vivo* screens) as well as a panel of published reference capsids from prior *in vivo* selections for CNS transduction (Chan *et al*., 2017; Deverman *et al*., 2016; Hanlon *et al*., 2019; Ravindra Kumar *et al*., 2020). The references included members of the AAV-PHP.B family known to utilize the LY6A receptor to cross the BBB (Hordeaux *et al*., 2019; Huang *et al*., 2019) (Supplementary file 8). Each 7-mer amino acid (AA) sequence in the library was encoded by two nucleotide sequences, which served as biological replicates.

**Figure 2.**
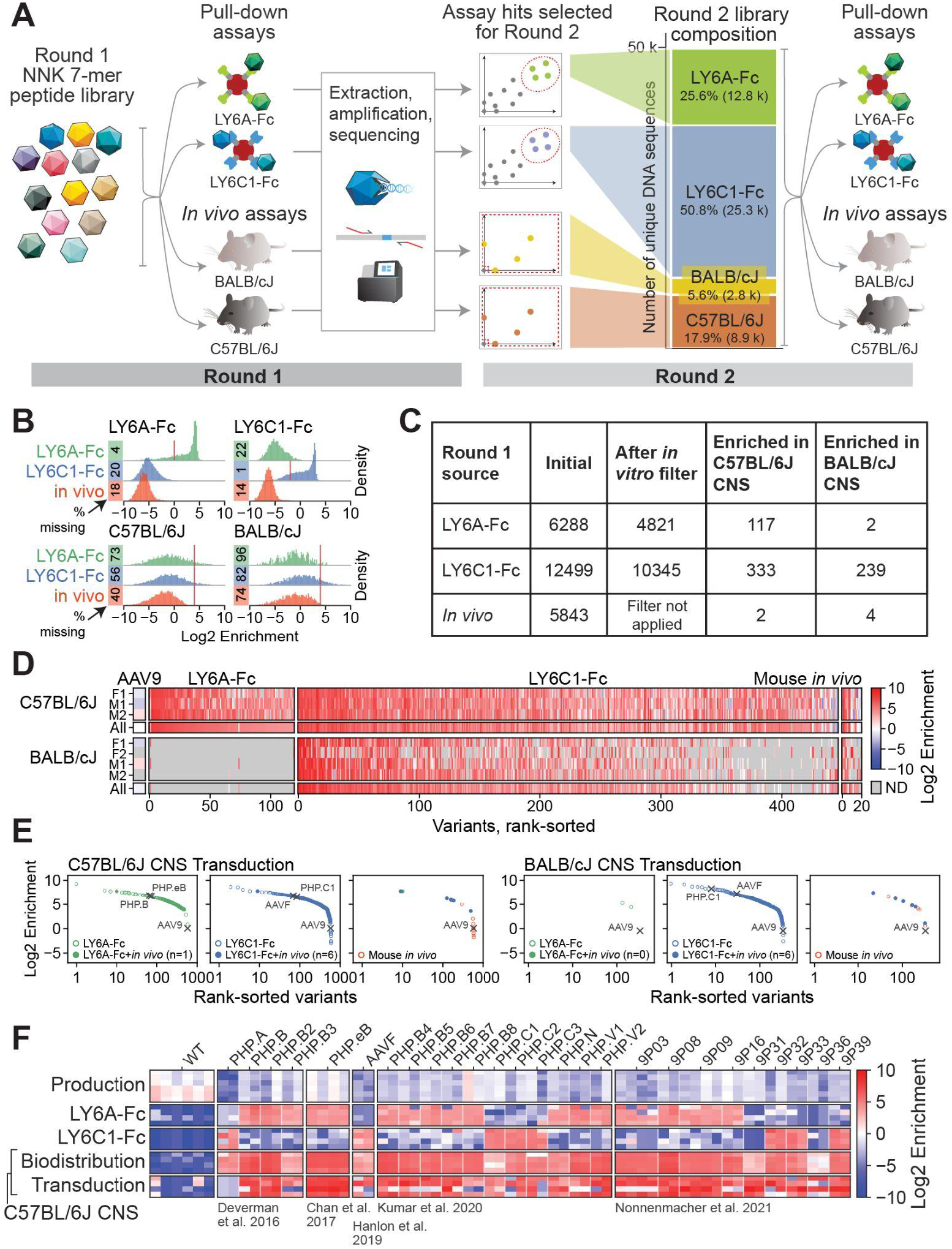
Round 2 validation of LY6A- and LY6C1-binding variants identified thousands of capsids with CNS transduction activity. (**A**) The Round 2 library was composed of a selection of top performing variants from the Round 1 assays for LY6A binding, LY6C1 binding, *in vivo* CNS transduction, and published reference sequences. The Round 2 library was subjected to screening as in Round 1. (**B**) The distributions show the Round 2 library performance in the pulldown assays and the CNS transduction of BALB/c and C57BL/6 mice, grouped and colored by the Round 1 selection source of each variant. The red lines indicate the thresholds set for the filters applied in (**C**). (**C**) The hits in Round 2 were identified as follows: target-binding capsids from the Round 1 screen were first filtered on their respective target binding activity in the Round 2 screen (LY6A: log_2_ enrichment > 0, LY6C1: log_2_ enrichment > −2). Variants were then filtered on Round 2 *in vivo* CNS transduction (log_2_ enrichment > 4 in either mouse strain and detected in at least 2 animals within that strain). (**D**) The *in vivo* log_2_ enrichment scores in C57BL/6J and BALB/cJ mice of Round 2 library variants that were filtered for high *in vivo* log_2_ enrichment scores in (**C**). The scores in individual animals (M*, F*) for each strain are shown alongside the average across animals (all). Variants are shown grouped and colored by their Round 1 selection source and rank-sorted on a combined score of C57BL/6J and BALB/cJ transduction. (**E**) The filtered variants from (**C**) are shown grouped and colored by their Round 1 selection source, and rank-sorted separately for each mouse strain. Reference controls and AAV9 are marked with crosses. Variants identified in both the Round 1 pull-down assays and *in vivo* screen are displayed as filled dots. (**F**) The target binding and C57BL/6J CNS biodistribution or transduction phenotypes of reference capsids with CNS tropisms are shown. Each capsid is represented by at least two 7-mer AA replicates (each column indicates a separate replicate).

The Round 2 library was screened *in vitro* and *in vivo* as in Round 1 (Figure 2A). The Round 2 data showed high agreement between replicates both *in vitro* and *in vivo* (Figure 2—figure supplement 1) and between 7-mer AA replicates across all assays (Figure 2—figure supplement 2). The majority of sequences identified in the Round 1 pull-down assays were validated by selective binding to their expected target in the Round 2 library screens (Figure 2B). When assessed for their ability to transduce cells in the CNS of C57BL/6J or BALB/cJ mice, hundreds of the capsid sequences identified by the pull-down assays with LY6A or LY6C1 were highly enriched (log_2_ enrichment > 4; Figure 2B,C). In comparison, far fewer sequences identified in the Round 1 *in vivo* screen were enriched in the Round 2 screen (Figure 2B,C). As previously observed for the AAV-PHP.B family, LY6A-binding capsids were highly enriched for *in vivo* transduction in the brains of C57BL/6 mice, but not BALB/cJ mice (Figure 2C,D). In contrast, numerous LY6C1-binding capsids were highly enriched in both mouse strains. These findings align with the expression levels of the two targets across strains (Huang *et al*., 2019).

We ranked the capsids identified by the pull-down assays and *in vivo* screening in Round 1 based on their enrichment in the CNS of C57BL/6J or BALB/cJ mice in Round 2 (Figure 2D). The ranking included previously characterized reference capsids such as AAV-PHP.B and AAV-F. Numerous capsids identified through the pull-down assays ranked among the top performing capsids in the *in vivo* selection alongside the reference capsids (Figure 2E). Notably, of 26 reference capsids identified in four prior independent studies using three different selection strategies in mice, 24 bound to LY6A or LY6C1 *in vitro;* 9P31 and 9P36 (Nonnenmacher *et al*., 2021) did not detectably bind to either LY6A or LY6C1 under our assay conditions (Figure 2F). These results suggest that LY6A and LY6C1 are capable of efficiently mediating the transportation of AAVs into the CNS, and that engineering capsids to bind proteins with such abilities can be an effective strategy to enhance tropism in a predictable manner based on target expression.

### Identification of a cluster of brain-enriched capsids from the Round 2 *in vivo* screen

The best performing CNS-transducing capsids in our Round 2 *in vivo* screen were LY6A or LY6C1 binders; however, we investigated the small subset of CNS-transducing capsids from the top Round 1 *in vivo* hits which did not bind to either target in the Round 2 pull-down assays (Figure 2—figure supplement 3A, log_2_ enrichment LY6A-Fc < 0, LY6C1-Fc < −2, C57BL/6J or BALB/cJ > 2, n = 180; these capsids did not pass the more stringent *in vivo* enrichment cutoff implemented in Figure 2C). Clustering of these capsids by pairwise hamming distance resulted in many small clusters and one larger cluster (Figure 2—figure supplement 3B). The large cluster had generally high BALB/cJ CNS transduction but lower efficiency in C57BL/6J, and exhibited a clear motif of *N*[T/V/I][R/K]** (Figure 2—figure supplement 3C,D, Supplementary file 9). Sequences in this cluster resemble that of our recently published AAV-BI30 capsid (AAV9 with the 7-mer insertion NNSTRGG), which highly transduces endothelial cells throughout the CNS of multiple mouse strains and rats *in vivo*, as well as human brain microvascular endothelial cells *in vitro* (Krolak *et al*., 2022).

### AAV capsids developed via pull-down assays effectively deliver genes to the mouse CNS

Top hits from the Round 2 *in vivo* selection were nominated for individual *in vivo* testing in BALB/cJ and C57BL/6J mice. First, we clustered the variants in the LY6A- and LY6C1-binding subsets from the Round 2 library that exhibited a log_2_ enrichment of more than 0 or −2, respectively (Figure 3A, Supplementary file 10). Not all LY6A-binding clusters yielded capsids that were enriched in the C57BL/6J brain in the Round 2 *in vivo* selection (Figure 3B). To test sequences in different clusters identified via the pull-down assays, five variants were selected for individual characterization based on their (1) mean brain transduction enrichment scores, (2) consistency of observed enrichment across replicates in the Round 2 screens (Figure 3C), (3) sequence diversity (the variants AAV-BI48, AAV-BI49, AAV-BI28, AAV-BI62, AAV-BI65 each represent different clusters as shown in Figure 3A), and (4) production fitness (estimated from the enrichment of the variants in the virus library compared to the plasmid library). Compared to AAV9, each variant exhibited enhanced CNS transduction consistent with their mechanism of action; AAV-PHP.eB, AAV-BI48 and AAV-BI49, which bind to LY6A, exhibited an enhanced CNS tropism in only C57BL/6J mice, whereas AAVF, AAV-BI28, AAV-BI62, and AAV-BI65, which bind to LY6C1, maintained their enhanced tropism across both mouse strains (Figure 3D).

**Figure 3.**
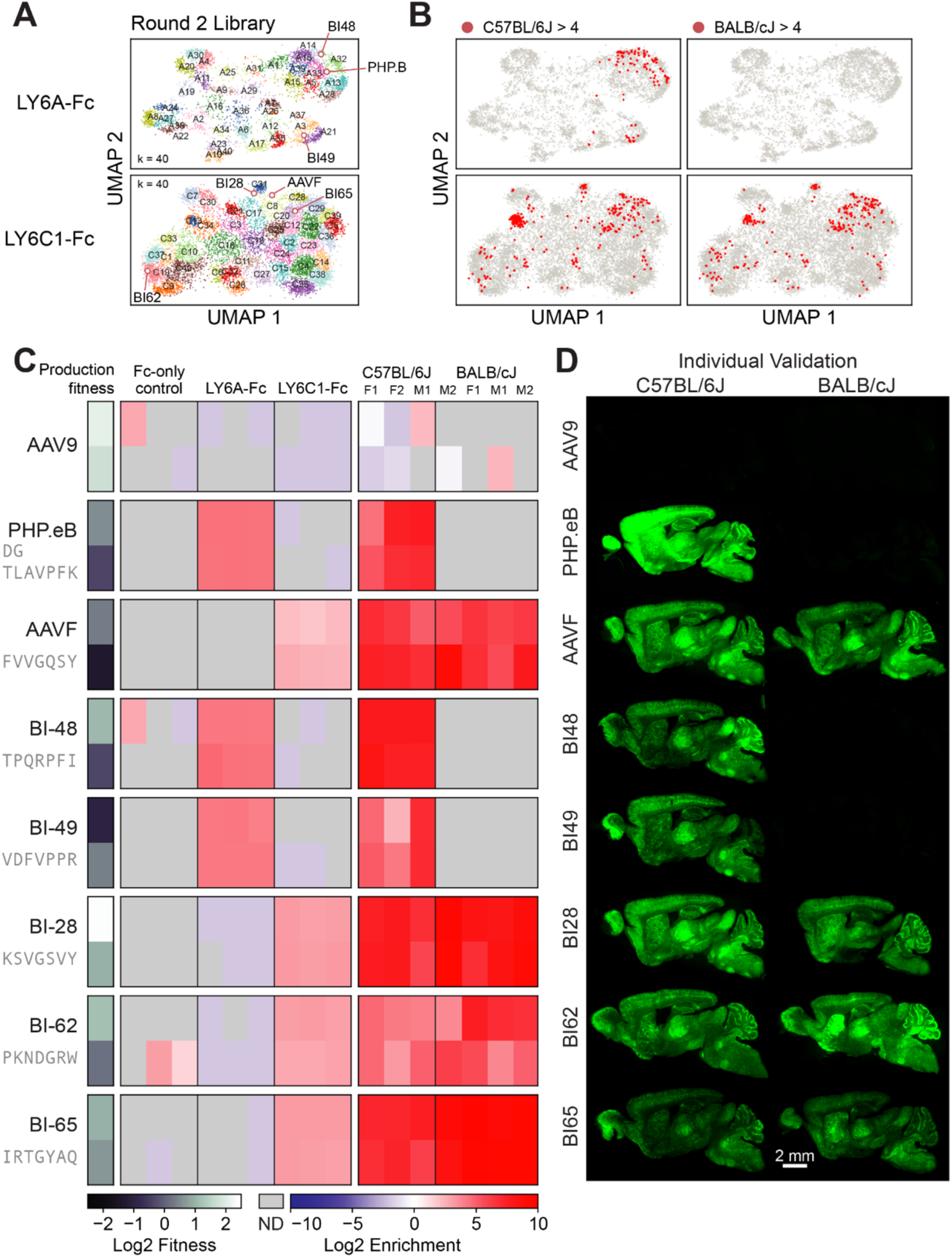
LY6A- and LY6C1-binding capsids identified in the pull-down assays cross the mouse BBB. (**A**) The UMAPs of Round 2 library variants are shown projected onto the UMAPs of Round 1 variants. Variant sequences were clustered with K-means (LY6A, k = 25; LY6C1, k = 30) (see cluster summaries in Supplementary file 10). (**B**) The Round 2 variants with an *in vivo* brain transduction log_2_ enrichment of > 4 in C57BL/6J mice (left) and BALB/cJ mice (right) are marked in red. (**C**) The Round 2 *in vivo* screen results for the reference capsids and five Round 2 variants selected for individual characterization are shown. Each variant is represented by two 7-mer AA replicates indicated by separate rows. ND = not detected. (**D**) Representative brain images are shown for the capsids in (**C**) that were individually tested in C57BL/6J mice (left) and BALB/cJ mice (right).

### The pull-down assay approach yields replicable, quantitative data that enables machine learning-guided sequence diversification

Screens for capsids with functions of interest typically only sample a small fraction of the theoretical sequence space (for a 7-mer insertion, the amino acid sequence space is 20^7^ or 1.28 billion). While it is impractical or even impossible (especially for longer sequences) to experimentally assay substantial portions of the sequence space, it is possible to train machine learning (ML) models using limited assay data to extend predictions to the rest of the unassayed sequence space. The highly replicable and quantitative pull-down assay data are amenable to ML-guided approaches for mapping 7-mer sequences to target binding.

To generate more diverse target-binding sequences, we sought to evaluate an ML-guided approach to train on data from only a single round of screening. We designed a library containing novel sequences generated using a supervised variational auto-encoder (SVAE) ML model or by saturation mutagenesis around specific motifs (Figure 4A; Figure 4—figure supplement 1A). As the Round 2 library was produced and assayed separately from the SVAE and saturation mutagenesis library, we used control sequences included in both libraries to perform calibration to account for the relative nature of enrichment and for batch effects (Figure 4—figure supplement 2). To generate variants via saturation mutagenesis, we chose to explore one highly enriched motif identified through LY6A and LY6C1 binding from the Round 1 screen: ***[K/R]PF[I/L] and ***G[W/Y]S[A/S], respectively (32K per motif; Figure 4A). These motifs were chosen as they were formed around residues with similar biochemical characteristics and contained many highly performant variants. The library containing the SVAE- and saturation mutagenesis-generated variants was subjected to pull-down assays and *in vivo* assays, and its results were compared to those of the Round 2 library.

**Figure 4.**
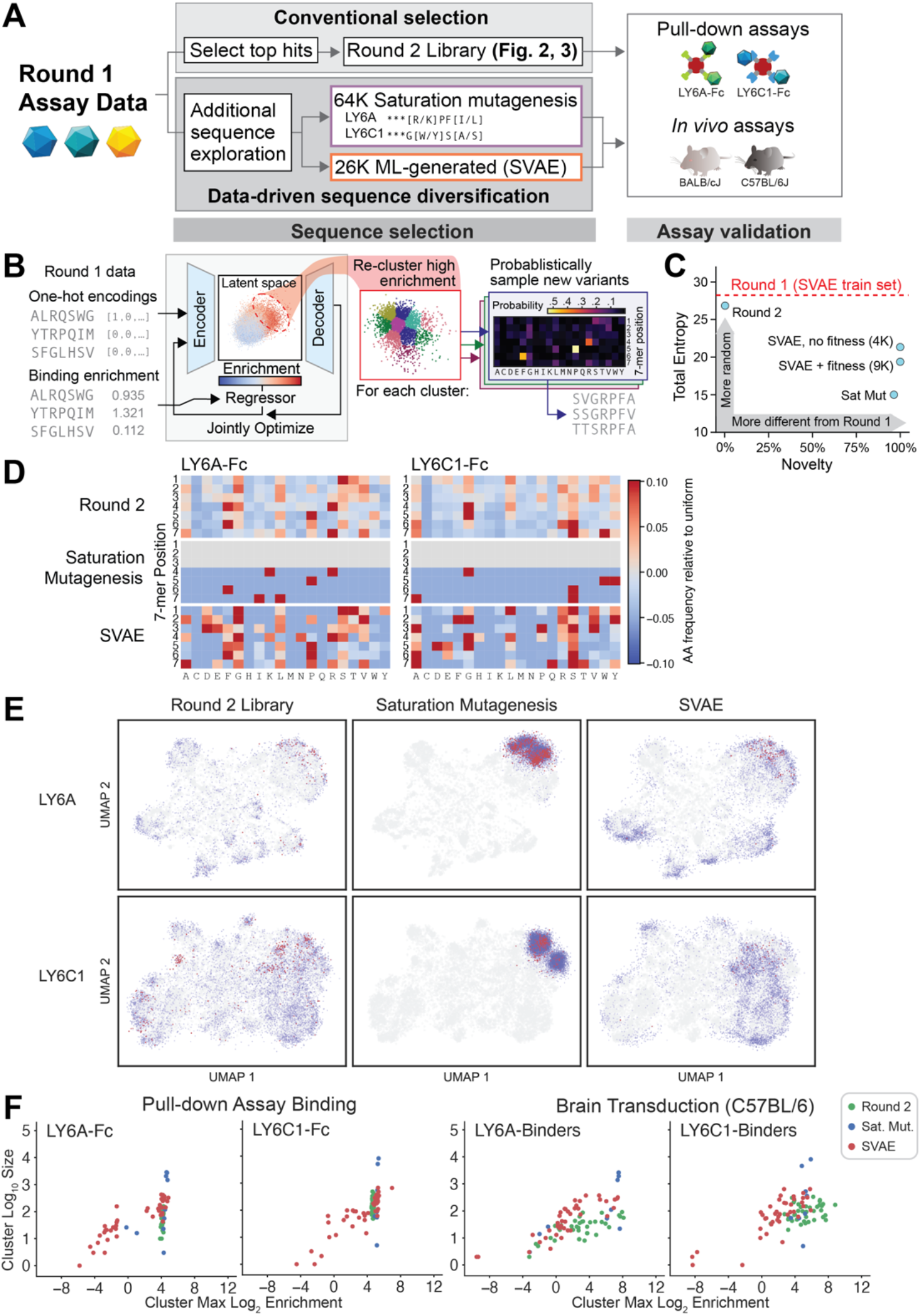
A single round of screening data can be used with supervised variational autoencoders (SVAE) and saturation mutagenesis to generate additional functional sequences. (**A**) Round 1 data were used to explore additional sequence diversity via two methods: saturation mutagenesis around two motifs (LY6A ***[K/R]PF[I/L], LY6C1 ***G[W/Y]S[A/S]) and SVAE ML generation. (**B**) The SVAE was trained on Round 1, library 1 sequences (encoder/decoder blocks) and binding enrichments (regression block). During training, these blocks were jointly optimized. High-binding enrichment sequences were isolated and re-clustered, and new sequences were sampled from each cluster’s probability weight matrix (PWM) (Figure 4—figure supplement 1 and Methods). (**C**) The total statistical entropy (summed entropies across all 7 amino acid positions) versus novelty (the fraction not found in Round 1) of each set of variants is shown. (**D**) Amino acid frequencies relative to uniform (1/20 chance of each) for the indicated libraries’ LY6A-Fc (left) and LY6C1-Fc binders (right). (**E**) The UMAP projection of sequence exploration for LY6A- (top row) and LY6C1-binders (bottom row) are mapped onto the same UMAP projection as Figures 1, 2, and 3; the Round 1 UMAP is reproduced in gray in each plot. Sequences with a log_2_ enrichment for production fitness > −1.0 (blue) and both fitness > −1.0 and *in vivo* log_2_ enrichment of > 3 (red) are shown for the Round 2 library (left), saturation mutagenesis (center), and SVAE (right). (**F**) Each point represents a cluster from (**E**), using the same cluster boundaries as in Figure 1F,G, plotted by cluster size versus the cluster’s maximum log_2_ enrichment in the binding or transduction assay. Log_2_ enrichments were calibrated using control sequences (Figure 4—figure supplement 2); no calibration adjustment exceeded 2.0.

To generate variants via ML, we used a sequence generation method based on latent-representation-learning models (Figure 4B; Figure 4—figure supplements 3 and 4), which have previously been applied toward the generation of a diverse set of viable capsids (Sinai *et al*., 2021). SVAE models for LY6A and LY6C1 binding were trained using one-hot encodings of 7-mer amino acid sequences and their associated target binding log_2_ enrichment. SVAE model accuracy was assessed by predicting binding enrichment on a held-out test set (Pearson correlation σ_A_ ≈ 0.83 and σ_c_ ≈ 0.85 for LY6A and LY6C1, respectively). Novel sequences with high predicted target binding were generated by clustering the high enrichment portion of the SVAE latent space, and then sampling from each cluster’s position weight matrix (amino acid frequencies of sequences in each cluster) (Figure 4B; Figure 4—figure supplements 3–5).

SVAE-generated variants were then evaluated *in silico* for predicted production fitness to preempt a high proportion of the variants failing to be produced at detectable levels in the virus library (Eid *et al*., unpublished). For the saturation mutagenesis approach, we generated all possible 7-mer sequences containing these motifs without filtering by predicted production fitness. The SVAE-generated variants were predicted to be prone to low production fitness (Figure 4—figure supplement 1B), likely as a result of optimizing solely for binding. Therefore, we generated two sets of variants via the SVAE: (1) 4K variants with highest predicted target binding (ignoring production fitness), and (2) 9K variants scoring the highest according to a joint score (Methods, Equation (3.1)) of predicted target binding and predicted production fitness (Figure 4—figure supplement 1B). After virus library production, ~25% of the 4K binding-only set was not detected, compared to <1% of the 9K joint score set (Figure 4—figure supplement 1C). Within the saturation mutagenesis-generated variants, 18.4% and 7.6% of the LY6A- and LY6C1-binding sets were not detected, respectively (Figure 4—figure supplement 1C).

Overall, the SVAE-generated variants were more diverse than the saturation mutagenesis-generated variants, which were designed around fixed motifs, as assessed by 7-mer entropy (Figure 4C), amino acid frequency (Figure 4D), and UMAP projection (Figure 4E). Both SVAE and saturation mutagenesis approaches generated top performers in *in vitro* target binding and *in vivo* brain transduction compared to the top hits from the Round 1 screen (Figure 4F; Figure 4—figure supplement 6); however, as anticipated, saturation mutagenesis based on a few select motifs produced variants which performed better on average than SVAE-generated variants that more extensively sampled the sequence space. These results demonstrate the inherent tradeoff between these two approaches, which are both viable sequence diversification strategies. The SVAE approach can explore more of the sequence space but produces less performant variants on average. In contrast, saturation mutagenesis can more comprehensively explore the space around a few high performing motifs to identify more hits possessing those motifs. Ultimately, the pull-down assays produced data from a single round of screening that could train ML models that capture sufficient understanding about the relationship between amino acid sequences and target binding performance.

## Discussion

We present a rapid method for enhancing the tropism of AAV vectors by introducing *de novo* interactions with proteins expressed on target cells. Our approach generated BBB-crossing capsids by first screening for direct *in vitro* interactions with specific host proteins rather than by selecting immediately for *in vivo* success. This mechanism-focused strategy identified thousands of capsids that specifically bind to the mouse brain endothelial cell surface proteins, LY6A or LY6C1, and many of these capsids exhibited enhanced CNS tropisms when validated *in vivo*, both in a pooled library and when tested individually. Importantly, the tropisms observed with LY6A- and LY6C1-binding capsids in different mouse strains matched expectations based on the strain-specific expression of these proteins. These data provide examples of how new virus capsid-receptor interactions can be introduced through the addition of short linear insertions into AAV capsid proteins.

*In vivo* selections typically recover a sparse subset of sequences with potential enhancements imparted through unknown mechanisms, which could be specific to a particular host strain or species and therefore not amenable to translational studies. In contrast, we demonstrate that a single round of protein target binding screening yielded highly reproducible and quantitative data based on a known mechanism of action. We leveraged these high-quality data to conduct saturation mutagenesis and ML-guided exploration of a more diverse target-binding sequence space to nominate additional, novel candidates for subsequent screening. Many of these novel candidates were found to exhibit high levels of *in vivo* CNS transduction in the validation library, again, within only two rounds of screening. Our findings demonstrate that saturation mutagenesis and ML-guided approaches both proved useful – with saturation mutagenesis comprehensively exploring the diversity around one or a few defined sequence motifs and SVAEs serving to explore a wider set of sequences – when used prior to the identification of the top functioning motifs.

The introduction of predictive modeling to target-specific capsid selection opens the capsid engineering process to a wide range of different computational approaches, with much more room for function-specific optimization and parameterization. A number of groups have generated diverse libraries of capsids for testing using ML-based approaches (Bryant *et al*., 2021; Shin *et al*., 2021), including unsupervised VAEs (Sinai *et al*., 2021). While our SVAE uses a standard one-hot encoding scheme which worked well using the high-quality data from the pull-down assays, others have experimented with the use of other encoding schemes such as physicochemical parameters (Georgiev, 2009) or learned representations (Alley *et al*., 2019). As the ML field and its role in biological research continues to develop, *in vitro* screening methods, such as the protein target binding assay used here, that generate high-quality, quantitative data will become crucial to take advantage of increasingly sophisticated computational approaches.

The method of engineering capsids through protein target binding assays enables screening against a wide range of host cell proteins. However, one limitation of this approach is that the features necessary for a protein to function as a new AAV receptor are not well elucidated. Valuable traits of these targets may include cell surface expression levels, intracellular trafficking routes, and a propensity for transcytosis in the case of interactions that mediate the crossing of vascular barriers. Identification of suitable protein targets is aided by recent advances in cell type characterization using single-cell transcriptomics and proteomics, e.g., the mouse endothelial atlas (Kalucka *et al*., 2020) and the human brain endothelial atlas (Garcia *et al*., 2022; Winkler *et al*., 2022). As more of the underlying biology of transcytosis and CNS transduction are characterized, mechanism-focused strategies offer a promising avenue for accelerating the development of capsids that bind defined CNS targets. An additional challenge in deploying this approach is the expression of less tractable host membrane proteins (e.g., proteins with multiple transmembrane domains) in the screening system. However, future improvements in complex protein expression should expand the range of host proteins that can be targeted in these screens. The pull-down assay approach demonstrated in this work should enable capsid library screening across protein targets from multiple host species to improve translation, as well as screening across different protein targets to identify capsids with highly specific target binding.

## Materials and Methods

### Capsid library cloning

The RNA expression system for the selection of functional AAV capsids was used as previously described (Krolak *et al*., 2022) with a modification to include a Woodchuck Hepatitis Virus (WHV) Posttranscriptional Regulatory Element (WPRE) between the restriction enzyme site Sall and Hindlll. The wild type AAV9 capsid gene sequence was synthesized (GenScript) with nucleotide changes at S448 (TCA to TCT, silent mutation), K449R (AAG to AGA), and G594 (GGC to GGT, silent mutation) to introduce Xbal and Agel restriction enzyme recognition sites for library fragment cloning.

For generating 7-mer NNK libraries, the hand-mixed primer Assembly-NNK-AAV9-588 (CCCGGAAGTATTCCTTGGTTTTGAACCCAACCGGTCTGCGCCTGTGCMNNMNNMNNMNNMNNMNNMNN TTGGGCACTCTGGTGGTT) encoding a 7-mer insertion between amino acid residues 588 and 589 of AAV9 was used as the reverse primer along with the Assembly-XbaI-F oligo (CACTCATCGACCAATACTTGTACTATCTCT) as a forward primer in a PCR reaction using Q5® High-Fidelity 2X Master Mix (NEB #M0492S) following the manufacturer’s protocol for 30 cycles with 10 ng pUC57-wtAAV9-X/A plasmid.

To assemble an oligonucleotide Library Synthesis (OLS) Pool (oligo pool; Agilent) into an AAV genome, 5 pM of the OLS pool was used as an initial reverse primer along with 0.5 μM Assembly-XbaI-F oligo as the forward primer to amplify and extend 10 ng pUC57-wtAAV9-X/A for five cycles. Then, the reaction was spiked with 0.5 μM of primer Assembly_AgeI-R (GTATTCCTTGGTTTTGAACCCAACCG) and amplified for an additional 25 cycles. The PCR product was purified using a Zymoclean DNA Gel Recovery kit (Zymo Research #D4007) following the manufacturer’s protocol. The 7-mer NNK or oligo pool PCR products were assembled into the RNA expression plasmid as previously described (Deverman *et al*., 2016).

### Virus production and titering

For both library and individual recombinant AAVs, viruses were generated by triple transfection of HEK293T/17 cells (ATCC, CRL-11268) using polyethylenimine (PEI), purified by ultracentrifugation over iodixanol gradients, and titered as previously described (Deverman *et al*., 2016; Krolak *et al*., 2022).

### Fc-fusion cloning and protein purification

The open reading frames of LY6A (NM_001271416.1) and LY6C1 (NM_010741.3) were separately cloned into an expression vector backbone with a C-terminal Fc-tag (Addgene plasmid #115773) using XbaI/EcoRV. The Fc construct DNA was transfected into HEK293T/17 cells (40 ug per 150 mm dish with PEI) in complete DMEM media with 5% FBS. 12-16 hours post-transfection, the plate was rinsed with PBS, and serum free media (Lonza, BEBP12-764Q) was added. Media containing the secreted Fc-fusion proteins was collected at 48 and 96 hours after the media change, filtered (Millipore SE1M003M00), and stored at 4°C until use. 35 μL of protein A-conjugated beads (ThermoFisher, 10001D) and Tween-20 (0.05% final concentration) were added to 30 mL of media and incubated at 4°C with end-to-end rotation. The next day, the beads were washed three times with DPBS containing 0.05% Tween-20. Expression was assessed by running a 5 μL aliquot of protein bound beads on a 4-12% protein gel; the remaining fraction was used for pull-down assay. In most cases, the beads were saturated with Fc-fusion protein as confirmed by the protein gel (data not shown).

### Pull-down assay

10 μL of Fc-fusion protein bound beads were mixed with 1e10 vg AAV capsid library in DPBS with 0.05% Tween-20 and 1% BSA and incubated overnight at 4°C. The next day, beads bound with virus were washed three times with PBS with 0.05% Tween-20, then treated with proteinase K to extract the viral genome which was purified with AMPure XP beads following the manufacturer’s protocol for PCR recovery and NGS sample preparation.

### Animals

All procedures were performed as approved by the Broad Institute IACUC (0213-06-08). BALBc (000651) and C57BL/6J mice (000664) were purchased from the Jackson Laboratory (JAX). Intravenous administration of rAAV vectors was performed by injecting the virus into the retro-orbital sinus.

### *In vivo* screening

For the selection in mice, 1e11 vg of the capsid libraries were intravenously injected into adult female animals. Two weeks post-injection, mice were euthanized and the brain and liver were collected. CNS transduction *in vivo* was measured by the transcribed capsid mRNAs as previously described (Krolak *et al*., 2022). Briefly, RNA was extracted from tissues with Trizol reagent followed by cleanup with RNeasy kit (Qiagen). 5 ug of RNA was converted to cDNA using Maxima H Minus Reverse Transcriptase (ThermoFisher, EP0751) according to manufacturer instructions and the resulting cDNA was used for capsid sequence recovery. In the Round 2 screen, one C57BL/6J mouse was excluded due to poor transduction by the virus library.

### Biodistribution and *in vivo* transduction of SVAE library

Eight-week-old C57BL/6 were injected intravenously with 1e11 vg of the SVAE virus library. For biodistribution, mice were anesthetized, perfused with PBS, and the brain and liver were collected two hours post-injection. Total DNA including the viral genome was extracted with a DNeasy kit (Qiagen) and used for NGS sample preparation. For *in vivo* transduction, tissues were harvested three weeks post-injection for RNA extraction and capsid sequence recovery with PCR.

### Tissue processing and imaging

Tissues were processed as previously described (Huang *et al*., 2019). Briefly, mice were anesthetized with Euthasol and transcardially perfused with PBS followed by 4% PFA in PBS. Sagittal brain sections were prepared with a vibratome (Leica). Images were taken with a Keyence BZ-X800 fluorescence microscope. All images were taken at the same magnification and exposure, and adjusted in photoshop with gamma = 1.4 and high = 150.

### NGS sample preparation

To prepare AAV libraries for sequencing, qPCR was performed on extracted AAV genomes or transcripts to determine the cycle thresholds for each sample type to prevent overamplification. Once cycle thresholds were determined, a first round PCR amplification using equal primer pairs (1-8) (PCR1 Primers) were used to attach Illumina Read 1 and Read 2 sequences using Q5 Hot Start High-Fidelity 2X Master Mix with an annealing temperature of 65°C for 20 seconds and an extension time of 1 minute. Round 1 PCR products were purified using AMPure XP beads following the manufacturer’s protocol and eluted in 25 μL UltraPure Water (ThermoScientific); then, 2 μL was used as input in a second round PCR amplification to attach Illumina adaptors and dual index primers (NEB, E7600S) for five PCR cycles using Q5 HotStart-High-Fidelity 2X Master Mix with an annealing temperature of 65°C for 20 seconds and an extension time of 1 minute. The second round PCR products were purified using AMPure XP beads following the manufacturer’s protocol and eluted in 25 μL UltraPure DNase/RNase-Free distilled water (ThermoScientific).

To quantify the amount of second round PCR product for NGS, an Agilent High Sensitivity DNA Kit (Agilent, 5067-4626) was used with an Agilent 2100 Bioanalyzer system. PCR products were then pooled and diluted to 2-4 nM in 10 mM Tris-HCl, pH 8.5 and sequenced on an Illumina NextSeq 550 following the manufacturer’s instructions using a NextSeq 500/550 Mid or High Output Kit (Illumina, 20024904 or 20024907), or on an Illumina NextSeq 1000 following the manufacturer’s instructions using NextSeq P2 v3 kits (Illumina, 20046812). Reads were allocated as follows: I1: 8, I2: 8, R1: 150, R2: 0.

### PCR1 primers

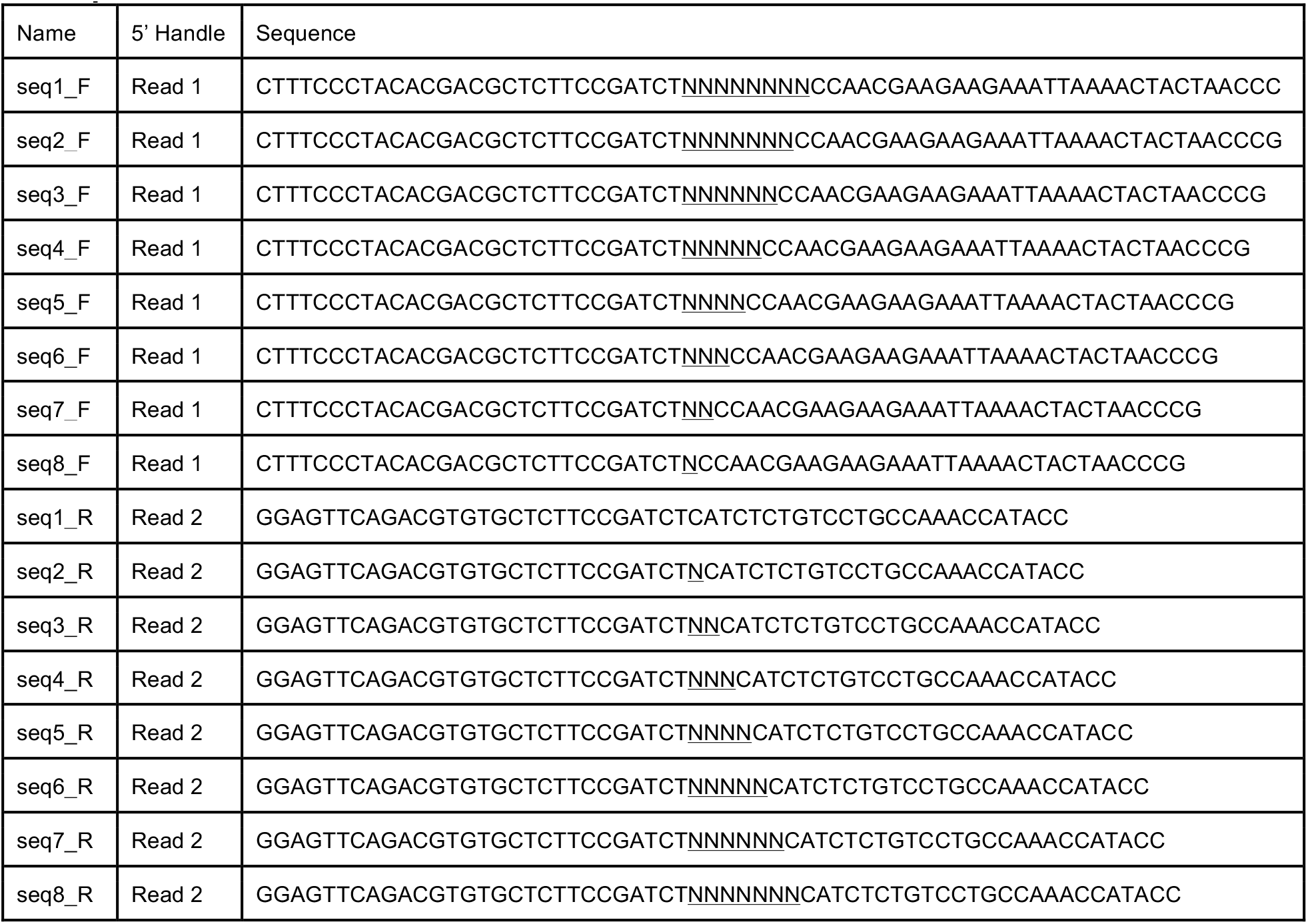

### NGS data processing

Sequencing data was demultiplexed with *bcl2fastq* (version v2.20.0.422) using the default parameters. The Read 1 sequence (excluding Illumina barcodes) was aligned to a short reference sequence of AAV9:

CCAACGAAGAAGAAATTAAAACTACTAACCCGGTAGCAACGGAGTCCTATGGACAAGTGGCCACAAACCA CCAGAGTGCCCAANNNNNNNNNNNNNNNNNNNNNGCACAGGCGCAGACCGGTTGGGTTCAAAACCAAG GAATACTTCCG

Alignment was performed with *bowtie2* (version 2.4.1) (Langmead and Salzberg, 2012) with the following parameters:

--end-to-end --very-sensitive --np 0 --n-ceil L,21,0.5 --xeq -N 1 --reorder --score-min L,−0.6,−0.6 −5 8 −3 8

The resulting sam files from *bowtie2* were sorted by read and compressed to bam files with *samtools* (version 1.11-2-g26d7c73, *htslib* version 1.11-9-g2264113) *(Danecek et al., 2021; Li et al., 2009)*.

*Python* (version 3.8.3) scripts and *pysam* (version 0.15.4) were used to flexibly extract the 21 nucleotide insertion from each amplicon read. Each read was assigned to one of the following bins: Failed, Invalid, or Valid. Failed reads were defined as reads that did not align to the reference sequence, or that had an in/del in the insertion region (i.e., 20 bases instead of 21 bases). Invalid reads were defined as reads whose 21 bases were successfully extracted, but matched any of the following conditions: 1) Any one base of the 21 bases had a quality score (AKA Phred score, QScore) below 20, i.e., error probability > 1/100, 2) Any one base was undetermined, i.e., “N”, 3) The 21 base sequence was not from the synthetic library (this case does not apply to NNK library), or 4) The 21 base sequence did not match a pattern, i.e., NNK (this case does not apply to the synthetic libraries). Valid reads were defined as reads that did not fit into either the Failed or Invalid bins. The Failed and Invalid reads were collected and analyzed for quality control purposes, and all subsequent analyses were performed on the Valid reads.

Count data for valid reads was aggregated per sequence, per sample, and was stored in a pivot table format, with nucleotide sequences on the rows, and samples (Illumina barcodes) on the columns. Sequences not detected in samples were assigned a count of 0.

To minimize the effect of sequencing error on analysis of the library data, variants in Library 1 and Library 2 with fewer than 10 total read counts across all samples and assays (including those not described in this paper) were excluded.

### Data Normalization

Count data was read-per-million (RPM) normalized to the sequencing depth of each sample *j*(Illumina barcode) with:

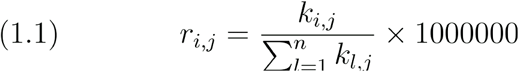

Where *r* is the RPM-normalized count, *k* is the raw count, *i* = 1, …, *n* sequences, and *j* = 1, …, *m* samples.

As each biological sample was run in triplicate, we aggregated data for each sample by taking the mean of the RPMs:

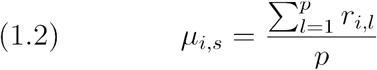

across *p* replicates of sample *s*. We estimated normalized variance across replicates by taking the coefficient of variation (CV):

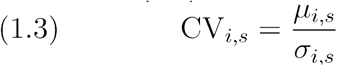

where *σ_i,s_* is the standard deviation for variant *i* in sample *s* over *p* replicates.

Log_2_ enrichment for each sequence was defined as:

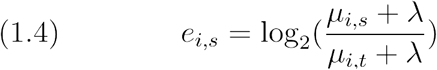

where λ is a pseudocount constant for ensuring valid values for the log transformation. For all data analyses, λ is set to 0.01.

### Clustering analysis

Target-specific capsids for LY6A and LY6C1 were selected according to their log_2_ enrichment for their respective receptor and the Fc-only control (blue highlighted regions in Figure 1D and Figure A—figure supplement 1A). The log_2_ enrichment cutoffs used for this analysis were: target > 8 when Fc enrichment missing (left marginal plot), target > 8 for Fc enrichment < = 0, target > (9/17 * Fc enrichment) + 8 for Fc enrichment > 0. This inclusion threshold yielded n = 5724 and n = 2291 LY6A-specific capsids for Library 1 and Library 2 respectively, and n = 7170 and n = 4214 LY6C1-specific capsids for Library 1 and Library 2 respectively. LY6A- and LY6C1-specific sequences were then separated from each other, and capsids from each target set were analyzed separately. Capsid sequences were one-hot encoded into vectors of length 20 x 7 = 140, and projected with UMAP with the following parameters: n_components = 2, n_neighbors = 200, min_dist = 0.15, metric = Euclidean. Capsid sequences were then clustered separately for LY6A and LY6C1, using their UMAP projection values (X1, X2) with the GaussianMixture model from scikit-learn (Pedregosa *et al*., 2011) with parameters n_components=40, random_state=1, n_init=10, max_iter=1000.

### Inter-library calibration

We produced and sequenced the Round 2 and combination SVAE/saturation mutagenesis libraries separately. The enrichment score of a variant in a library is derived from comparisons with the other members of the library, meaning that enrichment is a relative value. Thus, enrichments are comparable *within* a library, but not directly to other libraries. To enable comparisons between our two libraries, we included 3,352 variants in both for use in calibration of enrichment scores. We use a simple calibration method which adjusts enrichment scores to minimize the sum of errors between all shared variants.

Figure 4—figure supplement 2 shows the shared variants’ enrichments (A-C) and enrichment distribution after calibration (D). In addition to different library members, there is variation in the sequencing depth of the two libraries. Our calibration method does not account for sequencing depth, which we hypothesize causes some poorly enriched variants to show large enrichment differences between libraries (C, green box), so we chose to drop those variants from the calibration. Variants only detected in one library were also excluded from calibration. Note that while calibration improves comparisons between the libraries, error remains and can be substantial with standard deviations of 2.7, 2.1, and 2.3 for LY6A binding, LY6C1 binding, and the brain transduction assay respectively.

### Synthetic oligo pool library design and synthesis

The synthetic oligo pool library used for the secondary screening assay (Round 2) was obtained from Agilent. The oligonucleotides were designed to conform to the same template binding and assembly overlapping sequences as described above for the Round 1 NNK primers. The library oligo pool consisted of 7-mer insertion sequences recovered from the Round 1 pull-down assays based on the following criteria: (Library 1) Target log_2_ enr > 5, Target-Fc log_2_ enr - Fc control log_2_ enr > 3, and Target-Fc log_2_ enr - bead only control log_2_ enr >3 and were detected on at least 2 of the 3 replicates; (Library 2) Target log_2_ enr > 6 and filtration for specificity vs all other controls and assays based on Target RPM / SUM of all counts. The library also contained all of the top sequences recovered from the Round 1 C57BL/6J and BALB/cJ transcribed capsid sequence screen, published reference sequences and additional sequences screened for LY6A and LY6C1 binding through additional studies not described in this study. All sequences were encoded by two distinct nucleotide sequences designed to serve as biological replicates.

### Individual capsid characterization

Individual capsids were cloned into an iCAP-AAV9 (K449R) backbone (GenScript), produced as described above with a DNA genome that encodes nuclear localized GFP under a CAG promoter, and administered to C57BL/6J or BALB/cJ (Jackson Laboratory, 000664) mice at a dose of 3×10^11^ vg/mouse. Three weeks later, mice were perfused with 4% PFA. Tissue processing, immunohistochemistry, and imaging were performed as previously described (Huang *et al*., 2019).

### SVAE model

The data used in training the supervised variational auto-encoder (SVAE) were of the form

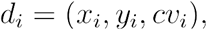

where *x_i_* is a one-hot encoded 7-mer AA sequence, *y_i_* the corresponding log2enr value (Eq 4) in the target assay, and *cυ_i_* the coefficient of variation (Eq 3). Only data points with assay mean RPM > 0 were included in training (at least 1/3 replicates had to be detected). The training/validation split was 0.8 and 0.2, respectively.

SVAE model architecture:

The SVAE (Figure 4—figure supplements 3 and 4) is composed of the following three neural network modules:

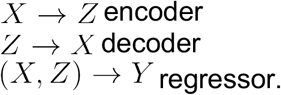

Among these, the encoder and decoder together form a standard VAE; the addition of the regressor enables supervision. The encoder learns a map

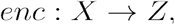

where Z is a latent space subject to the standard Gaussian prior (Kingma and Welling, 2013). The decoder learns a map

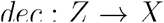

from the latent space back to the original (one-hot-encoded) sequence space. The regressor learns a map

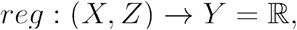

which takes as input a combined representation of a sequence *x_i_* and its (learned) latent representation *y_i_*, and maps it to a predicted log_2_ enrichment value 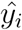.

In our model,

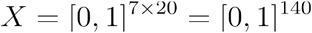

(such that 7-mer AA sequences are one-hot-encoded with respect to an alphabet of 20 amino acids),

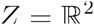

(i.e., we used a 2-dimensional latent space), and

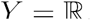

For our encoder, we used a 2-hidden-layer fully connected neural network with 100 and 40 nodes in the hidden layers, respectively, with ELU activation. Our decoder, constructed in mirror image of the encoder, was a 2-hidden-layer fully connected neural network with 40 and 100 nodes in the hidden layers, respectively. Our regressor was again a 2-hidden-layer fully connected neural network, but with 100 and 10 nodes in the hidden layers, respectively.

VAE Training:

The encoder and decoder networks are trained jointly with respect to the reconstruction loss

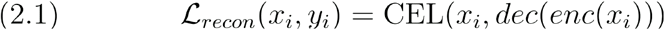

where CEL is the standard cross-entropy loss.

The regressor is trained with respect to the regression loss

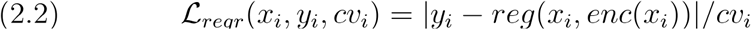

Additionally, there is a distributional loss term: 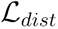 computed as the KL divergence of the VAE latent space and a standard gaussian prior (Kingma and Welling, 2013).

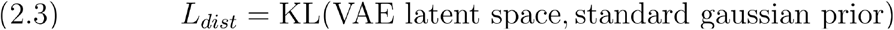

The overall loss of the SVAE is a linear combination of the 1) reconstruction loss, 2) regression loss, and 3) a distributional loss.

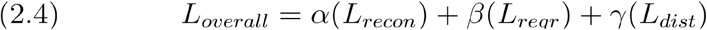

Where *α* = 1.0, *β* = 0.5, and *γ* = 0.1 parametrize the extent to which each loss term factors into the overall loss. These were tuned with hyperparameter optimization with the goal of producing a coherent latent space that separates the regressor values along a gradient.

Both models were trained until convergence, with a convergence threshold of 0.005. Convergence was identified as when the maximum difference between consecutive epochs across all loss metrics (*L_overall_*, *L_recon_*, *L_regr_*, and *L_dist_*) is less than the convergence threshold for 3 of 5 consecutive epochs. When trained according to this convergence criterion The LY6A-Fc and LY6C1-Fc model training ran for 64 and 63 epochs, respectively.

### SVAE Sequence Generation

After training, each model’s training data was projected into its trained 2D latent space. These points were clustered into 5 primary clusters using KMeans, using both latent space coordinates and log_2_ enrichment (Figure 4—figure supplement 5). The incorporation of the regression loss into training encourages points to separate spatially by enrichment value, along a gradient.

For each cluster, we calculated the mean enrichment of the sequences contained within it; because the latent space was encouraged to separate enrichment values along a gradient, and clustering was done using both latent space coordinates and enrichment value, the primary clusters formed clear high-, medium-, and low-mean-enrichment clusters (Figure 4—figure supplement 5). We isolated the single cluster with the highest mean enrichment to serve as the basis distribution for generating new sequences. This top cluster was then reclustered with K-means into 10 subclusters. Because the VAE’s latent space is trained to encode both sequence and corresponding assay enrichment, these subclusters correspond approximately to motif regions within the high performing cluster.

To generate new variants, for each subcluster, we encoded the amino acid frequencies at each position in the form of a position weight matrix (PWM). At each position, amino acids whose frequency was below the 80th percentile were filtered out. Using the remaining set of passing AAs per position, we generated all possible combinations of 7-mers. 7-mers already present in the training data were ignored.

### Optimization Library Composition

The Optimization Round 2 library consists of 96K capsid variants (two 7-mer AA replicates per variant, 192K DNA sequences total). This library of 96K variants comprises: 64K Saturation Mutagenesis, 26K SVAE-generated, 50 published/internal controls, 1K stop-codon controls, 6K calibration controls, and 4K positive training controls (Figure 4—figure supplement 1A). Within the same experimental pool as the Optimization Round 2 library were included an additional 26K sequences generated by an alternative VAE-based generation scheme. These additional sequences were used to compare the performance of the SVAE-based generation scheme as described in the text with that of the alternative scheme. We chose to present, in the comparison of library selection strategies (alongside saturation mutagenesis and standard selection), the scheme that generated the higher-performing set of variants on average.

The 26K SVAE-generated variants are equally divided between LY6A and LY6C1. Each receptor is further divided into two sets of sizes 4K and 9K, respectively, with different selection criteria: (1) the top 4K variants with the highest predicted binding enrichment according to the respective SVAE, and (2) the top 9K variants scoring the highest on a joint score depending on both high predicted binding enrichment and high predicted production fitness (Figure 4—figure supplement 1B). In order to compute the joint score across all novel generated variants per receptor, the set of predicted binding values is linearly scaled to lie in a range of [0,1]. The same is done for the fitness values. The scaled values are then simply added together (with equal weight) to compute the joint score. That is, if `gen_variants` is the full set of novel variants generated for either receptor, for a variant v in `gen_variants` the joint score `joint_score` (v) of v is defined as follows:

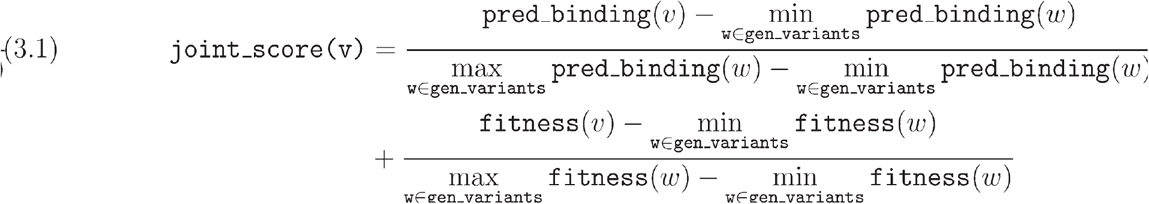

This split accounts for the SVAE’s lack of production fitness knowledge, i.e., the VAE model may not understand the destabilizing effects of certain AAs in the context of our 7-mer insert, such as cysteine (C) or tryptophan (W). The production fitness predictor is described in a separate article (Eid *et al*., unpublished).

The Top 4K subset, which was chosen on binding enrichment alone, displayed a markedly reduced observed production fitness, and a significant portion (24.3% for LY6A, 27.3% for LY6C1, Figure 4—figure supplement 1C) were not observed in the produced library—suggesting that these variants’ fitness was below our detection threshold. All subsequent analyses with SVAE-generated sequences use only the Top 9K subset.

The reference sequences (Supplementary file 11) include AAV capsids developed both in our lab and by other groups (Chan *et al*., 2017; Deverman *et al*., 2016; Nonnenmacher *et al*., 2021; Ravindra Kumar *et al*., 2020). 1K variants with stop codons in the 7-mer insert were included to assess the cross-packaging rate. 6K variants (3K each LY6A, LY6C1) are calibration controls, used to calibrate binding enrichment scores between this library and the training data (Round 1 Library 1). Each set of 3K variants was chosen to cover the dynamic range of each binding enrichment distribution. Finally, 2K variants were included as positive controls, 1K for each receptor, and sampled from the training data used to train each receptor’s respective SVAE model.

### Saturation Mutagenesis Library Generation

The Saturation Mutagenesis library consists of 8 motifs (4 motifs per receptor), with 8K variants per motif (64K total). Each motif has 4/7 positions fixed, leaving 3/7 flexible (20^3^ = 8,000 possible combinations). Starting from 236,951 sequences from our Round 1 library 1, we decompose each sequence into n-grams (motifs) of length 1–5. Wildcard positions within each motif are permitted with a maximum of 3 non-edge wildcards (e.g., A***A). The motif’s starting index (0-indexed) is appended to the end of the motif, to indicate motif position within the 7-mer, e.g., A<u>BCD</u>EFG → BCD1. Using this method, we build a bipartite graph of sequences on one side and motifs on the other, such that each sequence is linked to many motifs, and vice versa.

With this graph, we calculate several summary statistics for each motif: 1) motif “specific length”, the number of non-wildcard characters in the motif, e.g., A**A = 2; 2) the number of sequences linked to each motif; 3) a “motif enrichment”, the mean binding enrichments of the motif’s linked sequences. The specific length and number of sequences is useful for understanding motif specificity in the context of our Round 1 library 1. The more general the motif, the more its motif enrichment trends towards the population average of binding enrichments. For the Saturation Mutagenesis library, we chose motifs with a specific length small enough to admit thousands of variants per motif under combinatorial generation, but still with enough specificity to have a significant impact on enrichment. We selected a coherent set of motifs that exhibited high enrichment relative to other motifs of the same specificity.

The chosen motifs were PF4 for LY6A, and G*S3 for LY6C1. Given the constraints of our library size, we chose to select sub-motifs within these general motifs to fix for saturation mutagenesis. For LY6A, these were: ***KPFI, ***KPFL, ***RPFI, ***RPFL. For LY6C1, these were: ***GWSA, ***GWSS, ***GYSA, ***GYSS.

## Materials availability statement

All materials were acquired from commercial vendors as described. Packaging plasmids carrying the individually characterized LY6A or LY6C1 binding capsids will be made available by request and through Addgene.

## Data Availability

All code used in this study, as well as code and data for plot generation are available on GitHub: https://github.com/vector-engineering/AAVcapsidreceptor/

## Acknowledgements

We thank the members of the Deverman laboratory for continuous discussions of the project. This study was funded by the Stanley Foundation and Stanley Center for Psychiatric Research, the National Institute of Neurological Disorders and Stroke (UG3NS111689), Apertura Gene Therapy, and a Brain Initiative award funded through the National Institute of Mental Health (UG3MH120096) to B.E.D. Y.A.C. is supported by a Broad Ignite award. F.E.E. was supported by a Broad Shark Tank award.

## Author contributions

Conceptualization: BED, QH, KYC, FEE

Methodology: BED, QH, KYC, BA, ATC, HS, AB, FEE

Investigation: BED, QH, KYC, IGT, QZ, TB, BZ

Formal analysis: BED, QH, ATC, AB, HS, FEE, BA, YAC

Data curation: ATC, HS

Visualization: ATC, HS, AB, QH, YAC, BED

Funding acquisition: BED

Supervision: BED

Writing – original draft: QH, YAC, BED, ATC, HS, AB

Writing – review & editing: QH, YAC, BED, ATC, HS, AB

## Supplementary Files

Supplementary file 1. Library 1 UMAP clusters

Supplementary file 2. Library 1 LY6A UMAP cluster sequences

Supplementary file 3. Library 1 LY6C1 UMAP cluster sequences

Supplementary file 4. Library 2 UMAP clusters

Supplementary file 5. Library 2 LY6A UMAP cluster sequences

Supplementary file 6. Library 2 LY6C1 UMAP cluster sequences

Supplementary file 7. PHP.B-like UMAP clusters and sequences

Supplementary file 8. Round 2 Library reference sequences

Supplementary file 9. In vivo nonLY6A or LY6C1 binding sequences

Supplementary file 10. Round 2 Library UMAP clusters

Supplementary file 11. SVAE/Saturation Mutagenesis Library reference sequences

Supplementary file 12. Brain Transduction LY6A R1 top hits table

Supplementary file 13. Brain Transduction LY6C1 R1 top hits table

Supplementary file 14. Brain Transduction LY6C1 sat mut table

Supplementary file 15. Brain Transduction LY6C1 SVAE table

Supplementary file 16. LY6A R1 top hits table

Supplementary file 17. LY6A sat mut table

Supplementary file 18. LY6A SVAE table

Supplementary file 19. Brain Transduction LY6A sat mut table

Supplementary file 20. Brain Transduction LY6A SVAE table

Supplementary file 21. LY6C1 R1 top hits table

Supplementary file 22. LY6C1 sat mut table

Supplementary file 23. LY6C1 SVAE table

## Supplementary Figures

**Figure 1—figure supplement 1.**
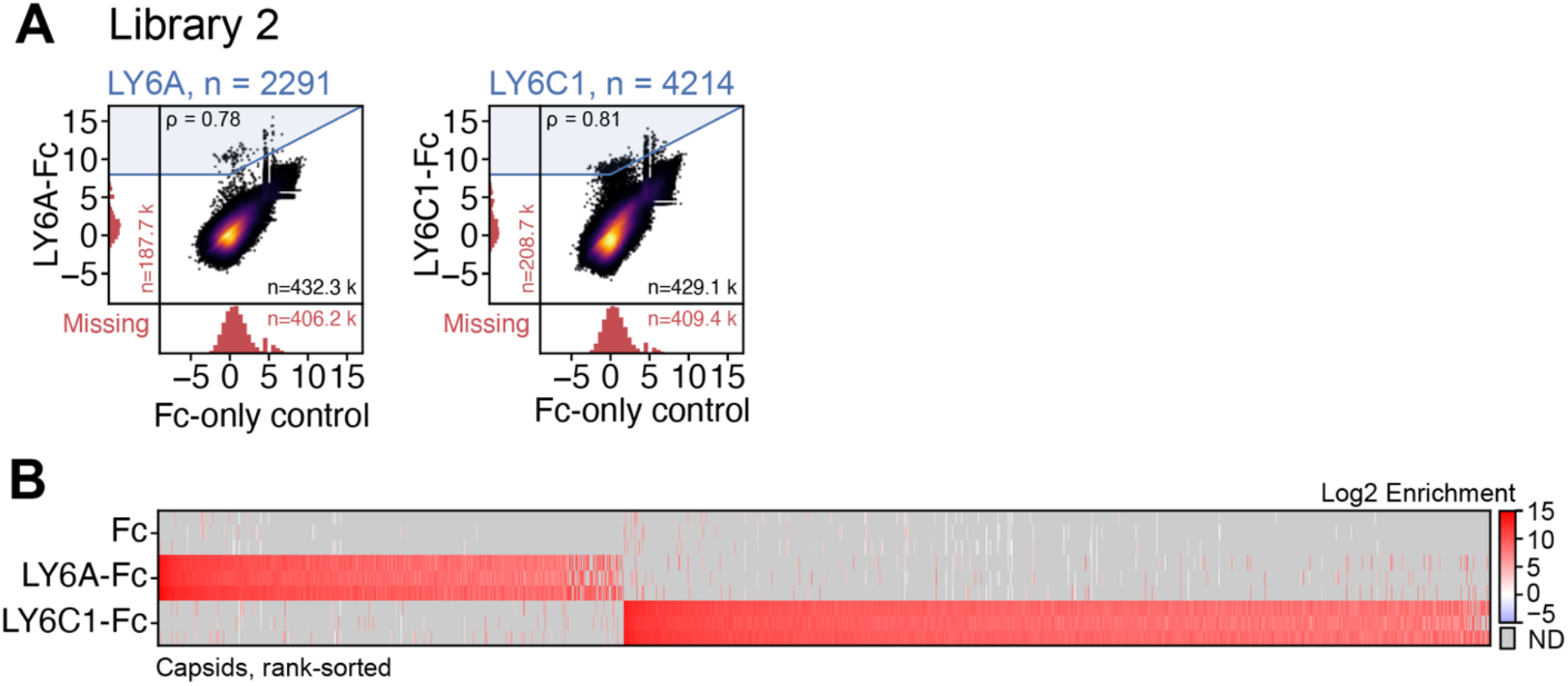
Identification of target-specific capsids using an independently generated random 7-mer library (Library 2). (**A**) The variant log_2_ enrichment (average RPM between replicates, normalized to the starting library RPM) plotted between LY6A-Fc or LY6C1-Fc versus the Fc-only control. The capsids detected in both assays are displayed in the upper-right quadrant. Missing variants from either assay are displayed in the marginal quadrants. (**B**) The log_2_ enrichment of selected variants highlighted in blue in (**A**) with each replicate’s enrichment plotted in separate rows (n = 3). ND = not detected.

**Figure 1—figure supplement 2.**
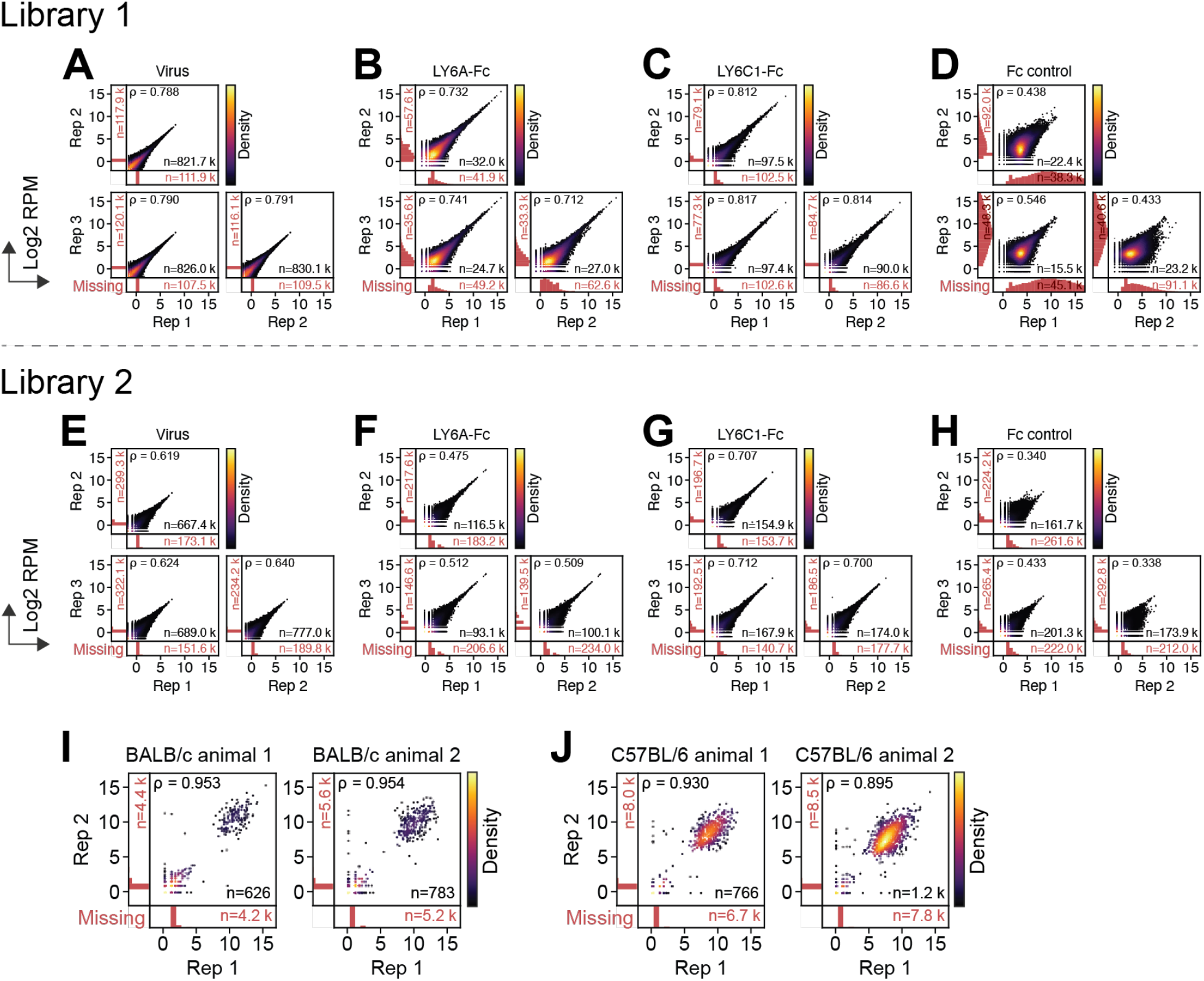
Replicability for *in vitro* binding assays and *in vivo* CNS transduction screens. (**A–D**) Screen of Library 1 replicability of the log_2_ RPM of the (**A**) starting virus library, (**B**) LY6A-Fc, (**C**) LY6C1-Fc, and (**D**) Fc-only control. (**E–H**) Screen of Library 2 replicability of the (**E**) starting virus library, (**F**) LY6A-Fc, (**G**) LY6C1-Fc, and (**H**) Fc-only control. (**I,J**) Replicability of separate RNA extractions (n = 2 extractions per mouse strain) within each mouse strain (n = 2 mice) for (**I**) BALB/cJ and (**J**) C57BL/6J. The capsids detected in both replicates are displayed in the upperright quadrant. The missing variants from either replicate are displayed in the marginal quadrants.

**Figure 1—figure supplement 3.**
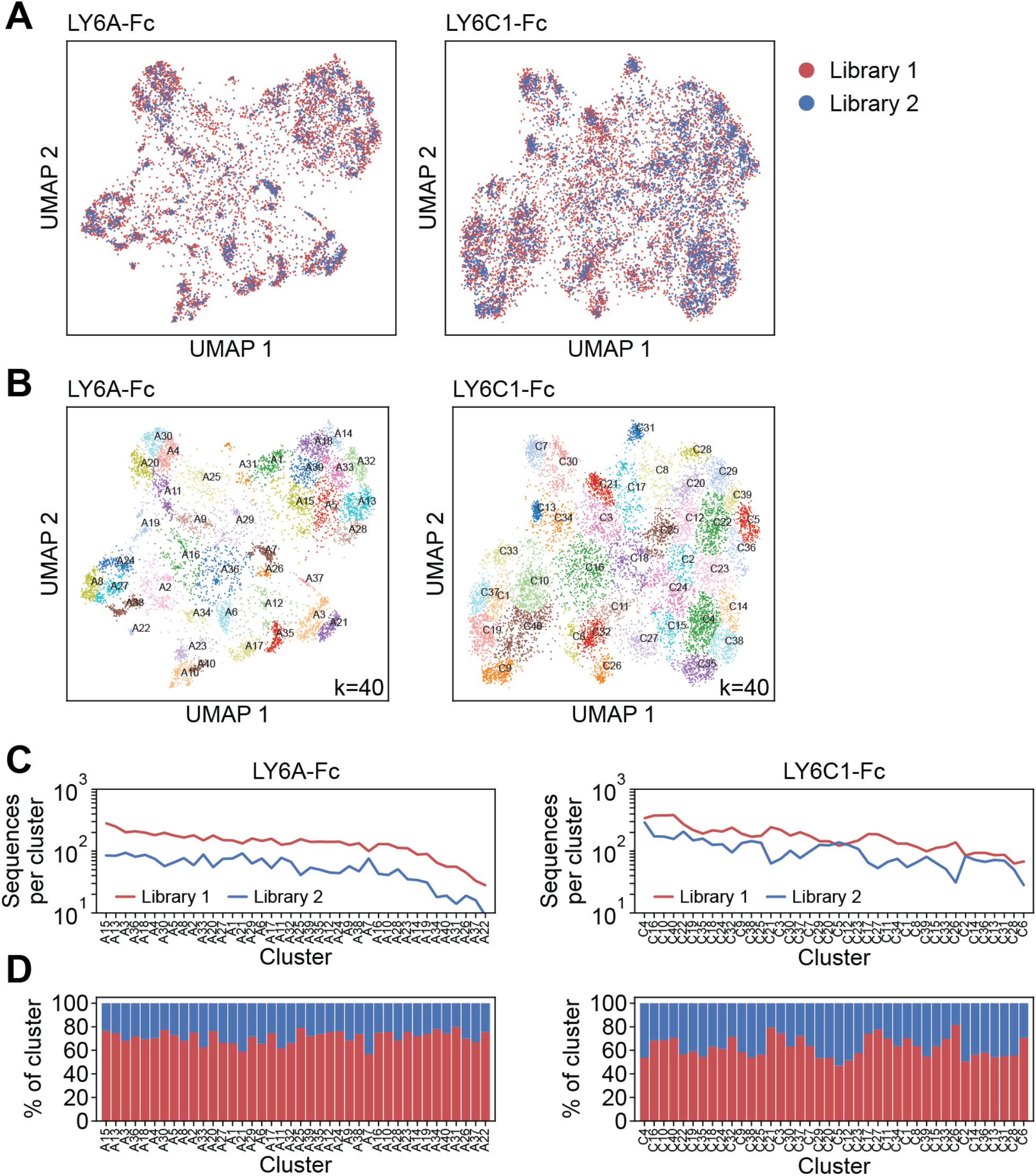
Joint clustering of *in vitro* binders identified from the two libraries. (**A**) The joint UMAP embedding of target-specific 7-mer sequences with sequences colored according to experiment. (**B**) The clustering (Gaussian mixture model, k = 40) on the joint embedding. (**C,D**) The number (**C**) and percentage (**D**) of 7-mer sequences by the Round 1 pull-down screen from each library, per cluster (sorted from left to right by the number of sequences per cluster).

**Figure 1—figure supplement 4.**
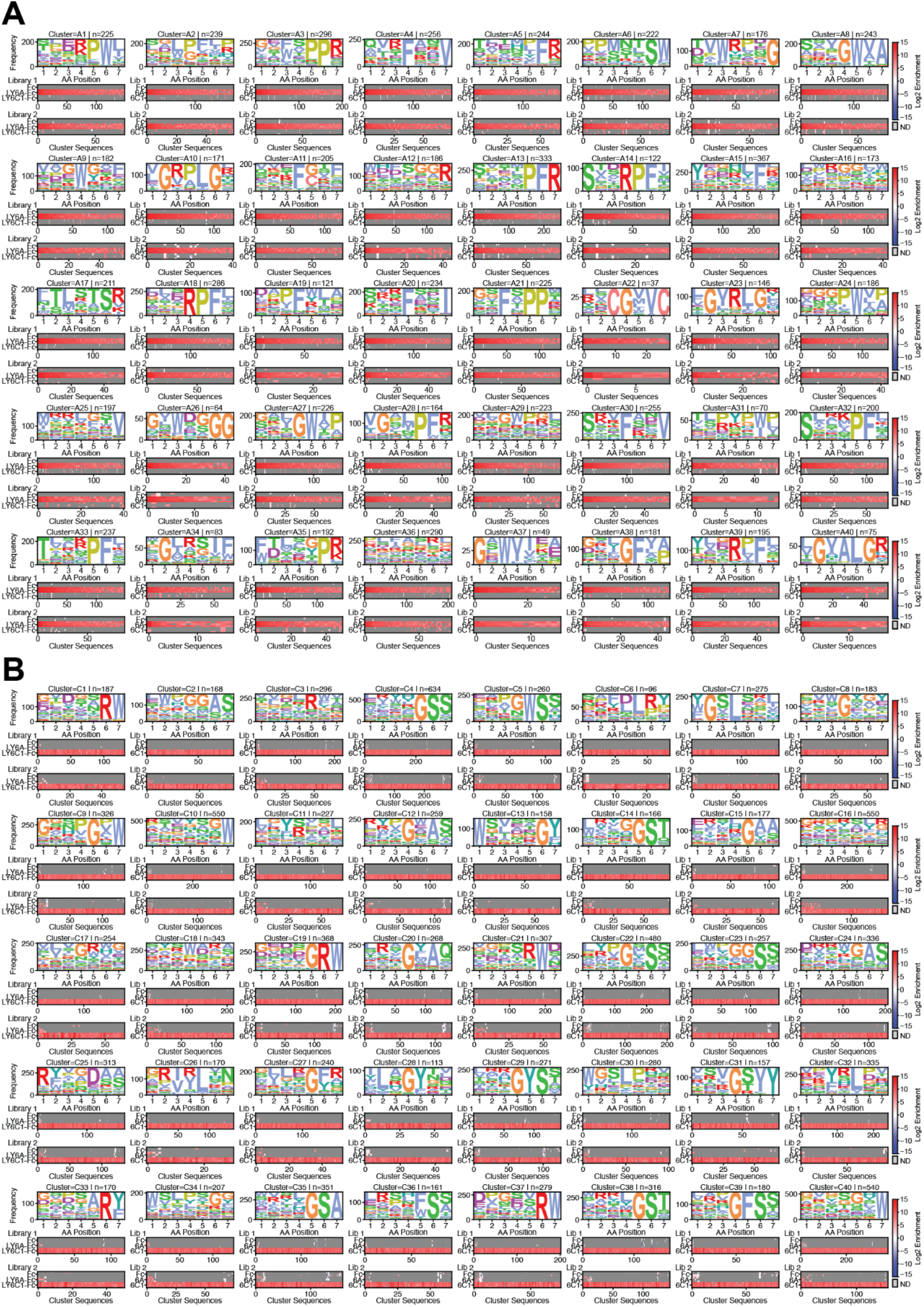
Cluster analysis of the Round 1 target-specific 7-mer sequences. (**A**) LY6A or (**B**) LY6C1 cluster sequence logos and the corresponding heatmap of log_2_ enrichments for sequences in each cluster for the Fc-only control, LY6A-Fc, and LY6C1-Fc.

**Figure 2—figure supplement 1.**
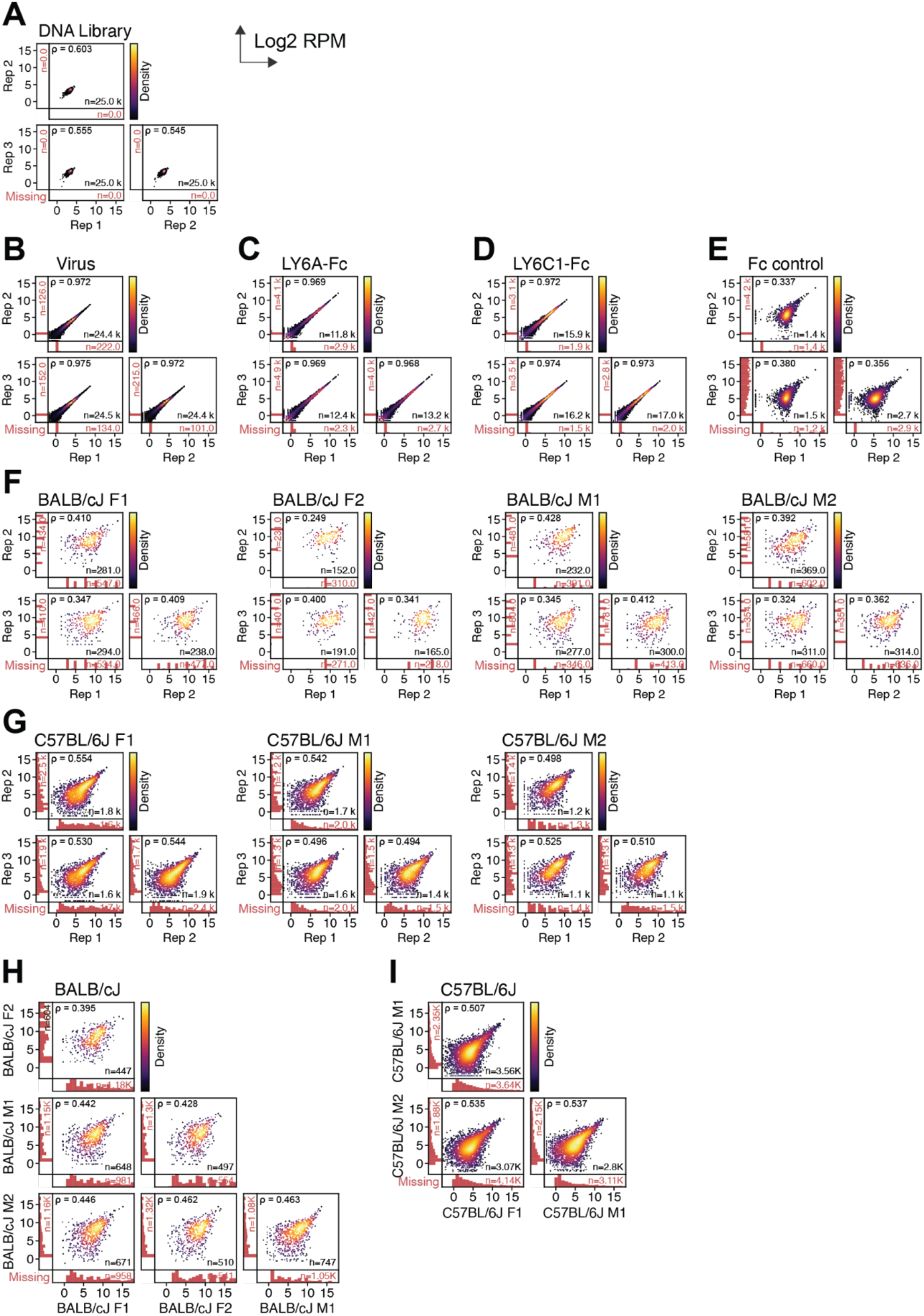
The Pearson correlations of Round 2 *in vitro* and *in vivo* replicates. The plots show the replicability of the log_2_ reads per million (RPM) of the (**A**) DNA (plasmid) library, (**B**) virus library, (**C**) LY6A-Fc, (**D**) LY6C1-Fc, and (**E**) Fc-only control. The capsids detected in both replicates are displayed in the upper-right quadrant. The missing variants from either replicate are displayed in the marginal quadrants. The replicability of separate RNA extractions are shown for (**F**) BALB/cJ (4 mice [F1, F2, M1, M2], n = 3 extraction replicates per animal) and (**G**) C57BL/6J (3 mice [F1, M1, M2], n = 3 extraction replicates per animal). The mean RPM from the extraction replicates between animals were compared for (**H**) BALB/cJ and (**I**) C57BL/6J.

**Figure 2—figure supplement 2.**
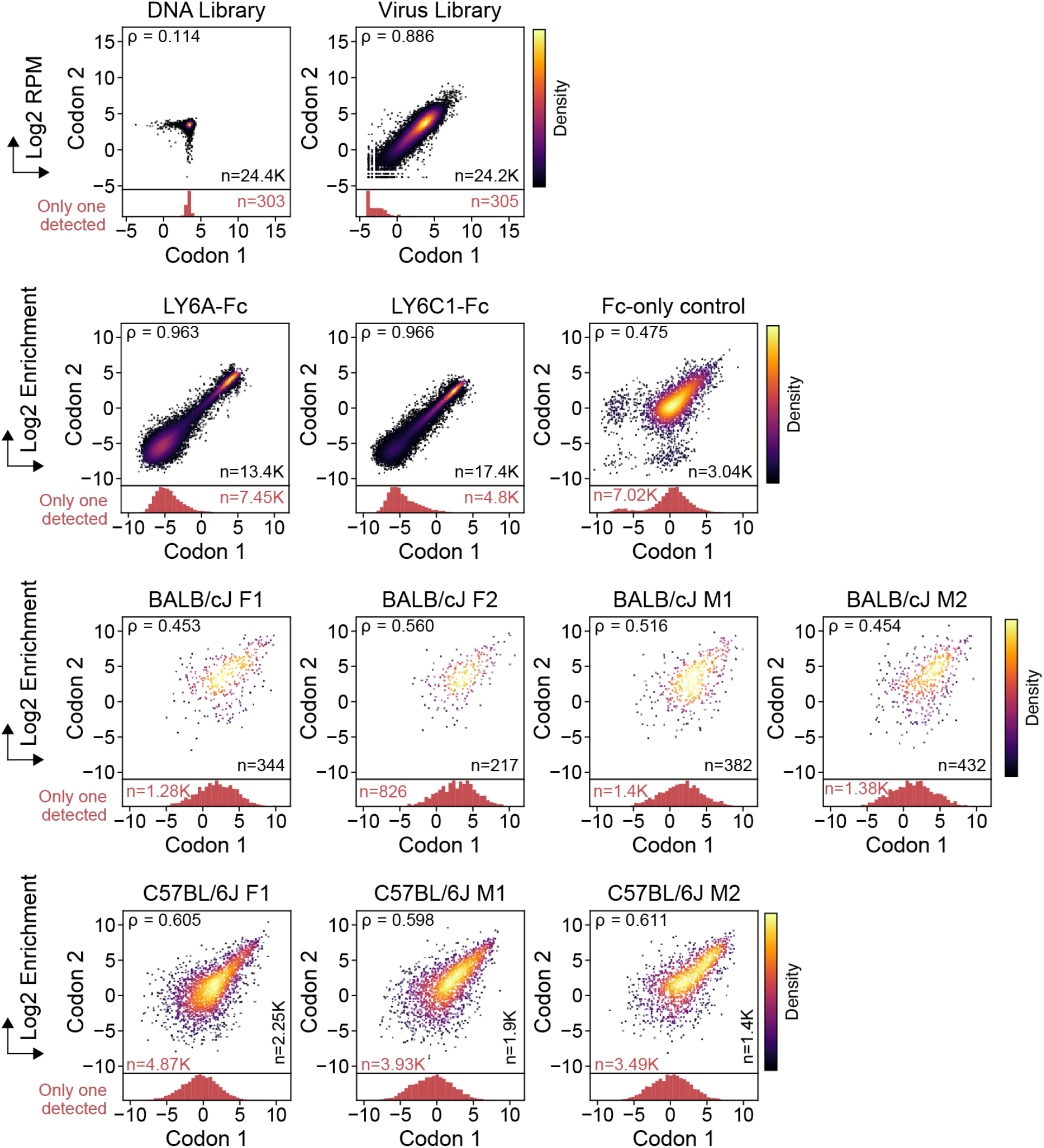
The Pearson correlations between AA replicates in the Round 2 library. The values shown are log_2_ RPM for the DNA library and virus library samples, and log_2_ enrichment for the *in vitro* and *in vivo* samples. Sequences within pairs of 7-mer AA replicates (codon 1 and codon 2) were randomly assigned to either the x- or the y-axis, with the exception of AA sequences missing their partner within a replicate pair that are assigned to the x-axis and plotted in the histogram below each plot.

**Figure 2—figure supplement 3.**
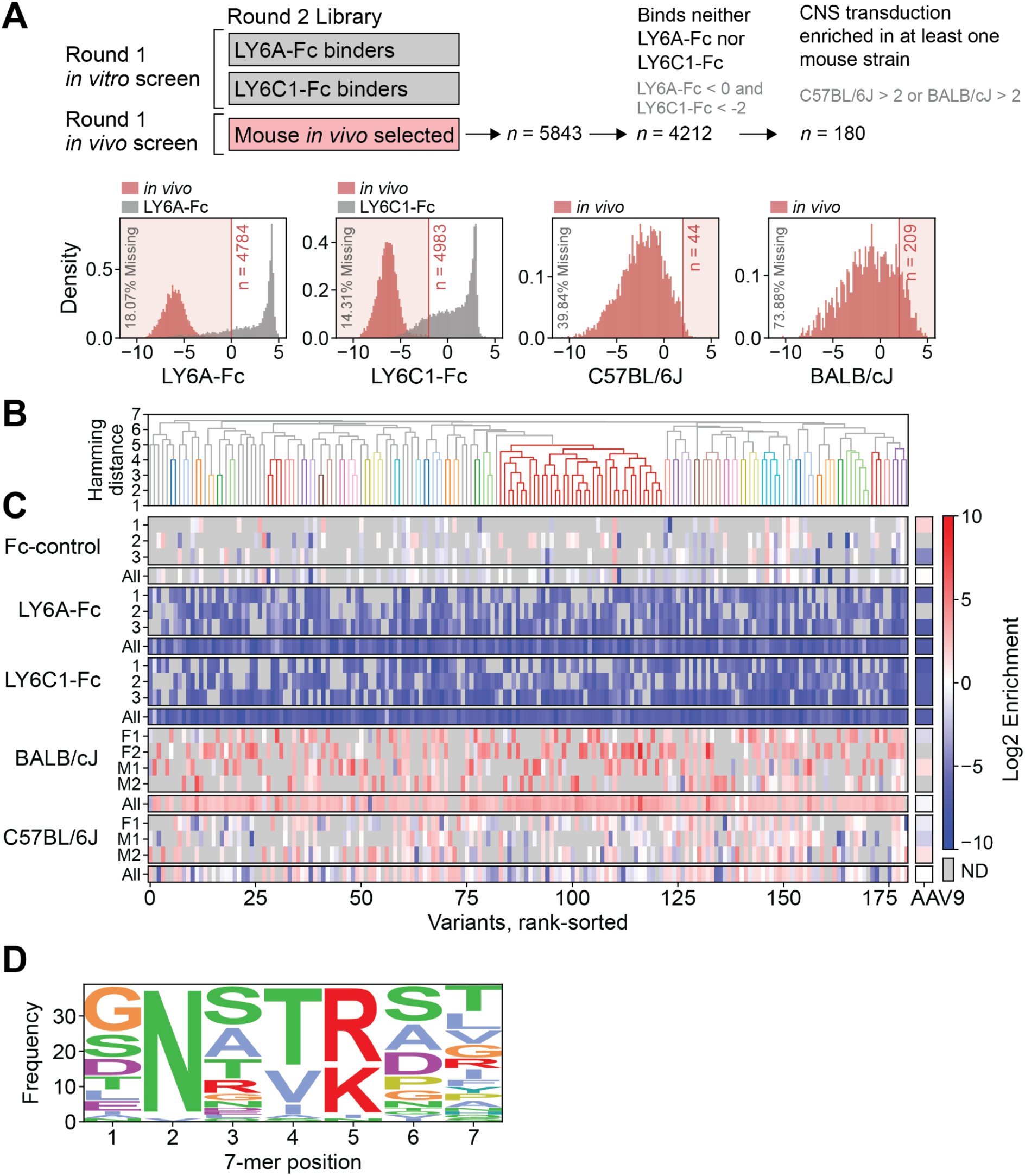
Identification of a brain-enriched motif that binds to neither LY6A-Fc nor LY6C1-Fc. (**A**) Round 2 variants identified in the Round 1 *in vivo* screen (red) were filtered by the thresholds shown for low binding to LY6A-Fc (LY6A-Fc binders are shown in gray), low binding to LY6C1-Fc (LY6C1-Fc binders are shown in gray), and high CNS transduction in either C57BL/6J or BALB/cJ mice. This combined filtering yielded 180 variants. (**B**) Hierarchical clustering of the 180 variants by hamming distance (linkage = average, cutoff = 5) yielded one large cluster (red, center, n = 39). (**C**) Log_2_ enrichment is shown for each variant ordered by the clustering tree in (**B**) for *in vitro* binding of the Fc-only control, LY6A-Fc, LY6C1-Fc, and CNS transduction in BALB/cJ or C57BL/6J mice. (**D**) The sequence motif of the center red cluster (n=39) in (**B**) shows a clear pattern of *N*[T/V/I][R/K]**.

**Figure 4—figure supplement 1.**
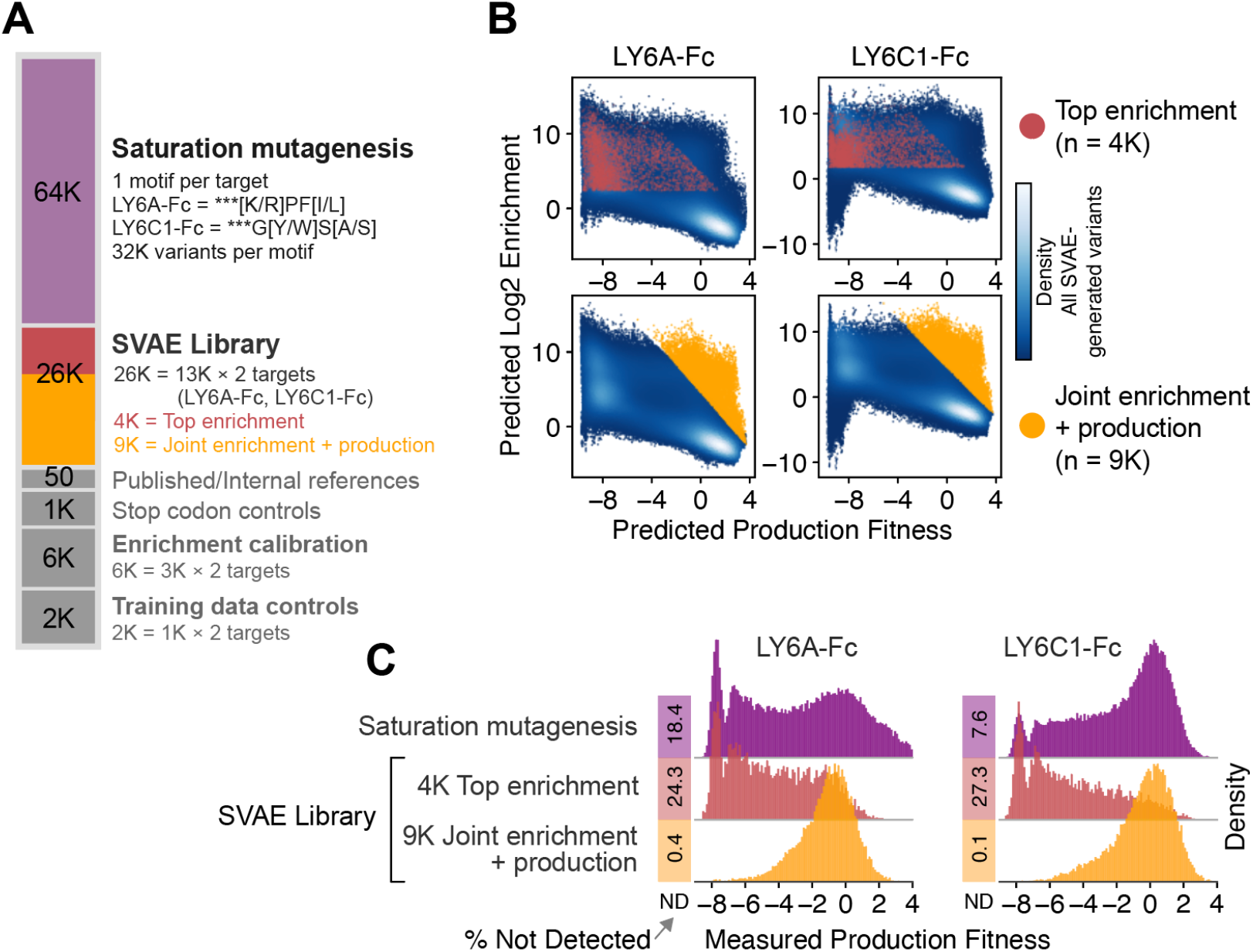
SVAE library composition and selection of SVAE-generated variants. (**A**) The combined SVAE and saturation mutagenesis library is composed of saturation mutagenesis variants generated from one motif per target (LY6A ***[K/R]PF[I/L], LY6C1 ***G[W/Y]S[A/S]) with 32K variants per motif; 13K SVAE-generated variants per target; 50 previously characterized variants from our group and the literature; 1K variants with stop codons to assess cross-packaging; 6K variants (3K per target) that were evenly selected across low-to-high enrichment bins to calibrate the enrichment scores from this library to the library used to train the SVAE models (Round 1, Library 1); 2K variants (1K for each target) that were randomly chosen from the SVAE training data (i.e., variants with non-zero RPM from Round 1) as training data controls. (**B**) The predicted binding enrichment and predicted production fitness for SVAE-generated variants (150K generated *in silico* per target) are shown. Included in the SVAE Library were the 4K variants with the top predicted binding enrichment according to the SVAE (red), as well as the top 9K variants according to a joint score of predicted binding enrichment and predicted production fitness (yellow). (**C**) The virus library shown in (**A**) was produced and the distributions of the measured production fitness of the saturation mutagenesis-generated and SVAE-generated variants are shown.

**Figure 4—figure supplement 2.**
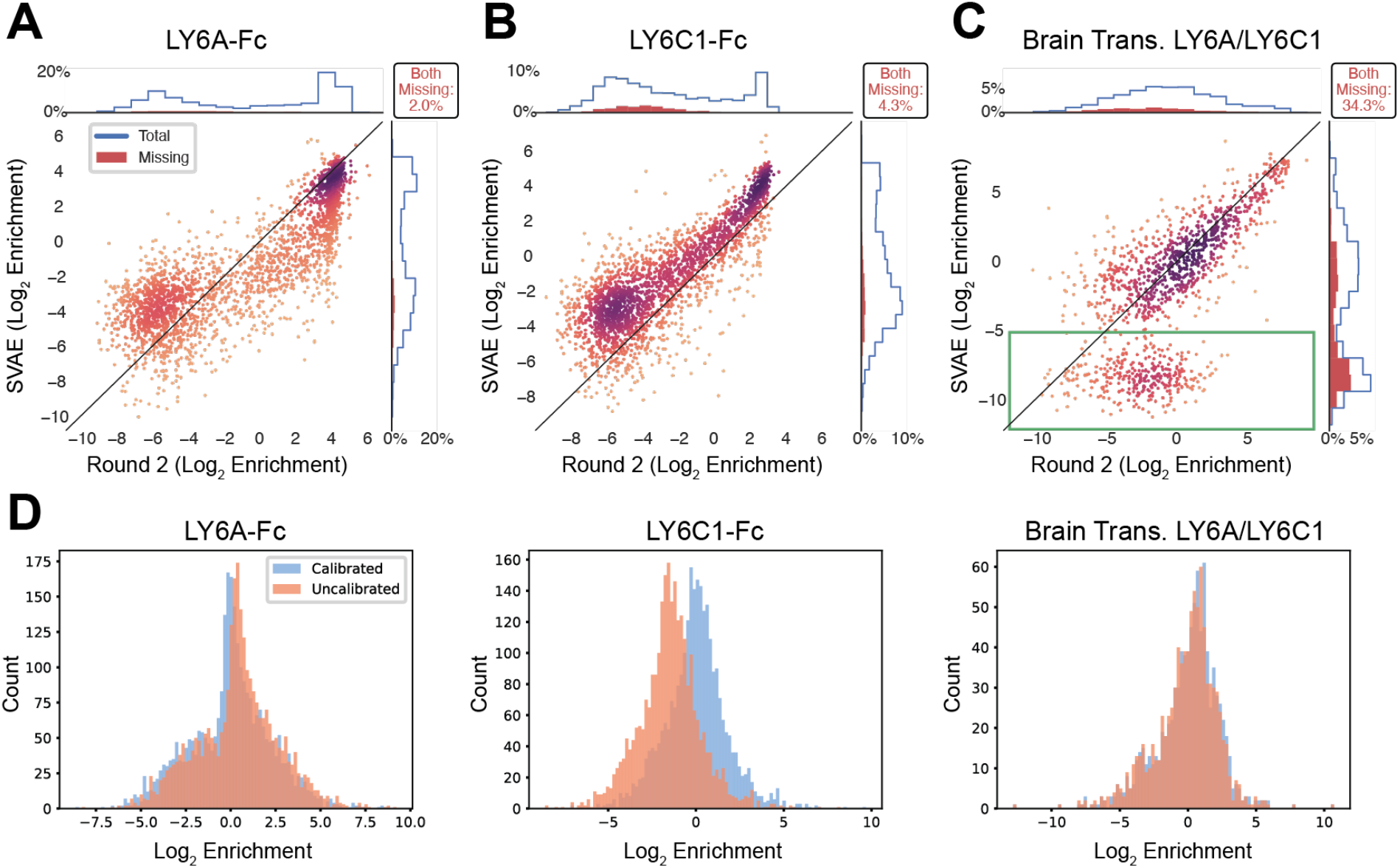
The calibration of the SVAE and saturation mutagenesis library to the Round 2 library. Library variant enrichment scores are relative because they are derived from comparisons to the other members of the same library. The scores of variants from separate libraries were calibrated by computing a single value that adjusts these scatter plots (**A–C**) on the y-axis to minimize error (see Methods, Supplementary files 12-23). (**A,B**) Plots of the enrichment scores within the SVAE library versus the Round 2 library for the uncalibrated LY6A- and LY6C1-binding sequences that are common to both libraries. Histograms on the top and right margins show the distribution of total variants (blue) and variants missing in one of the assays (red). (**C**) The same as in (**A,B**), but for the brain transduction assay. For the brain transduction assay, both libraries contain LY6A- and LY6C1-binding variants so a single calibration value was applied. Points in the green box were dropped when computing the calibration. We hypothesize that this discrepancy arose from the Round 2 library being sequenced more deeply. (**D**) Histograms of pre- and post-calibration enrichment for each assay. Calibration values are as follows: LY6A: −0.37, LY6C1: 1.50, brain transduction, combined LY6A/LY6C1: 0.14. The amount of shift between the pre- and post-calibration histograms corresponds to the calibration value for each assay.

**Figure 4—figure supplement 3.**
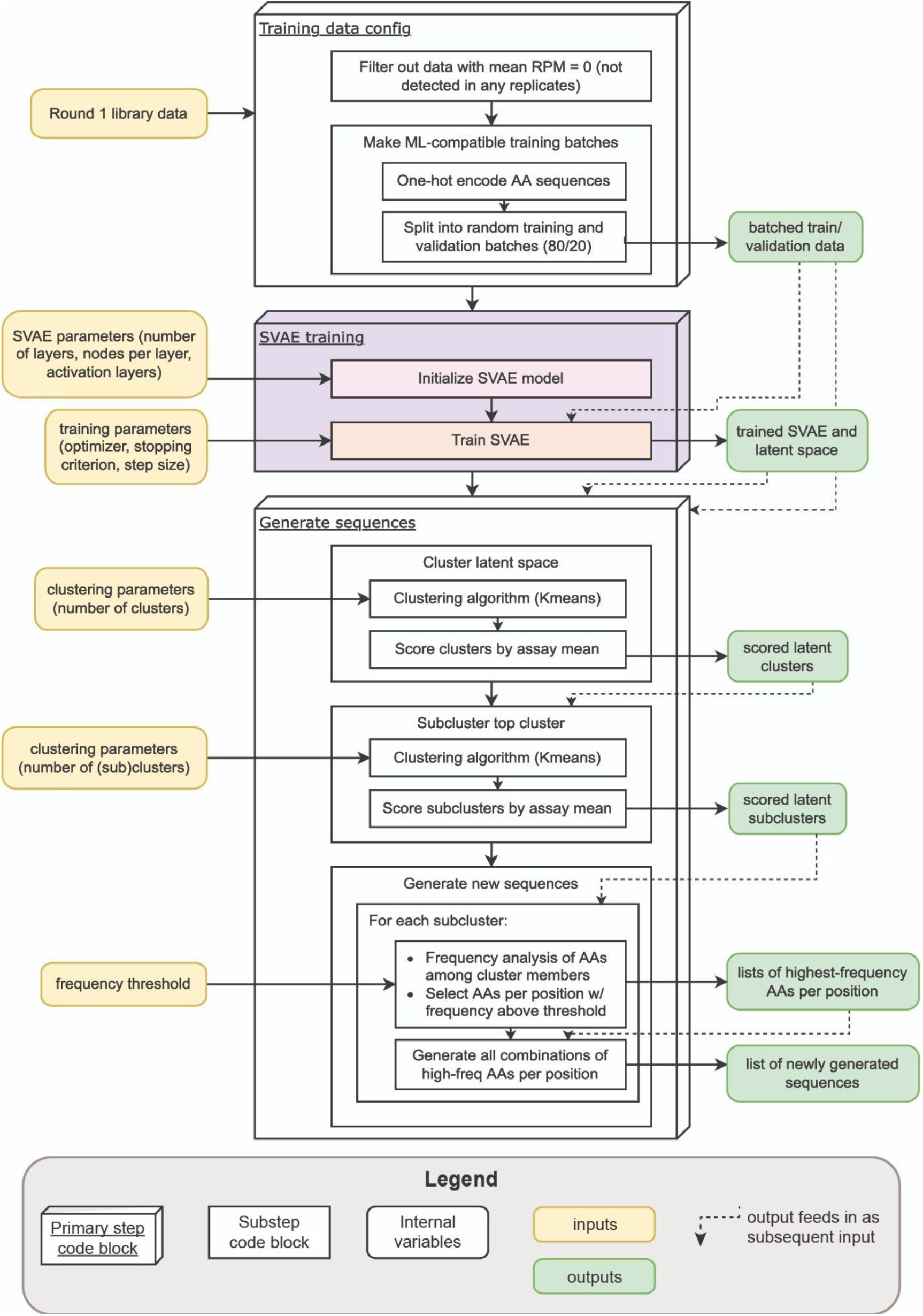
SVAE-based sequence generation procedure, SVAE model, and latent spaces. (**A**) A schematic of the complete SVAE-based sequence generation procedure, including (1) the processing of training data, (2) SVAE training, and (3) sequence generation using the SVAE latent space.

**Figure 4—figure supplement 4.**
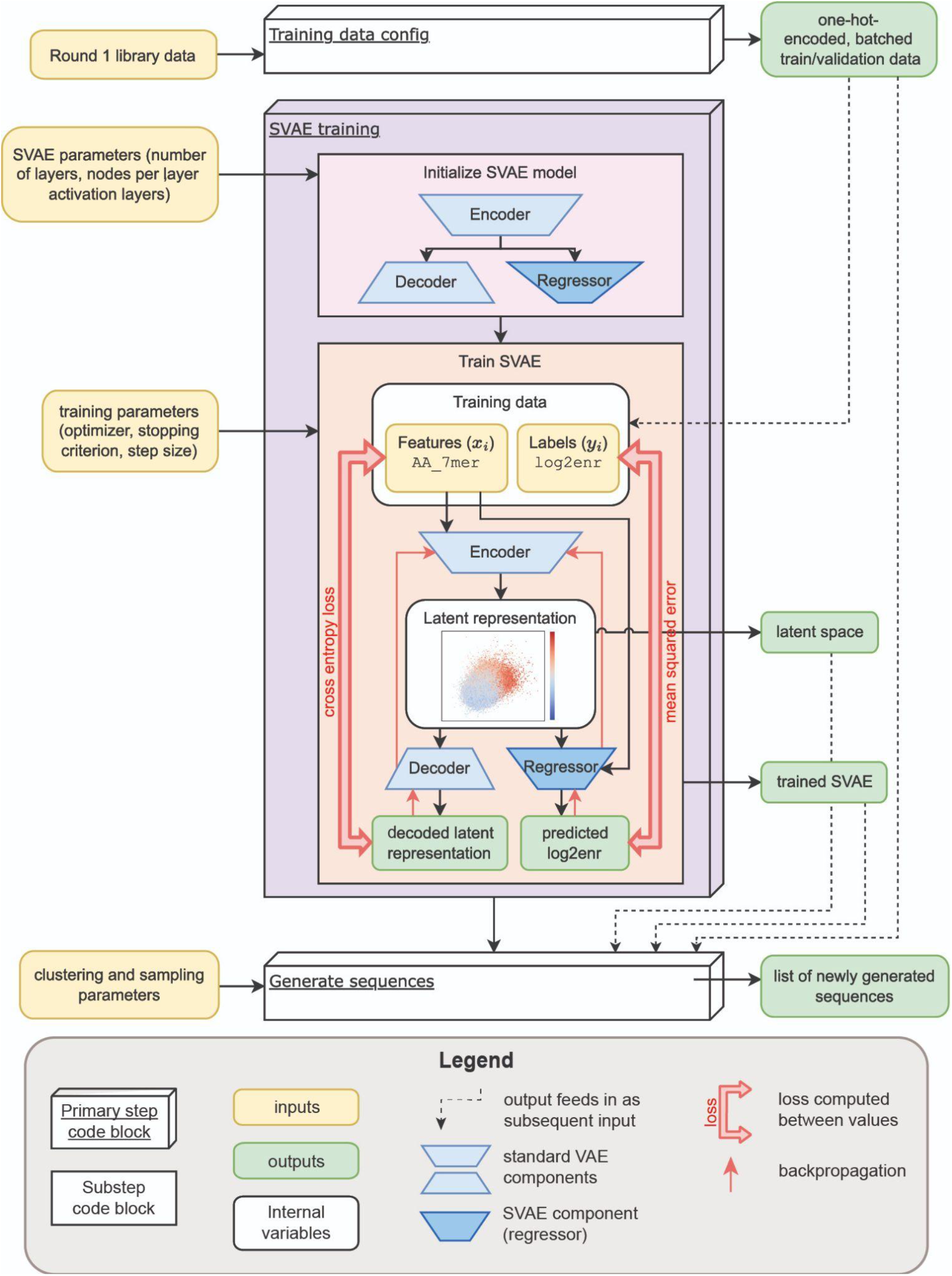
An expanded schematic of SVAE training.

**Figure 4—figure supplement 5.**
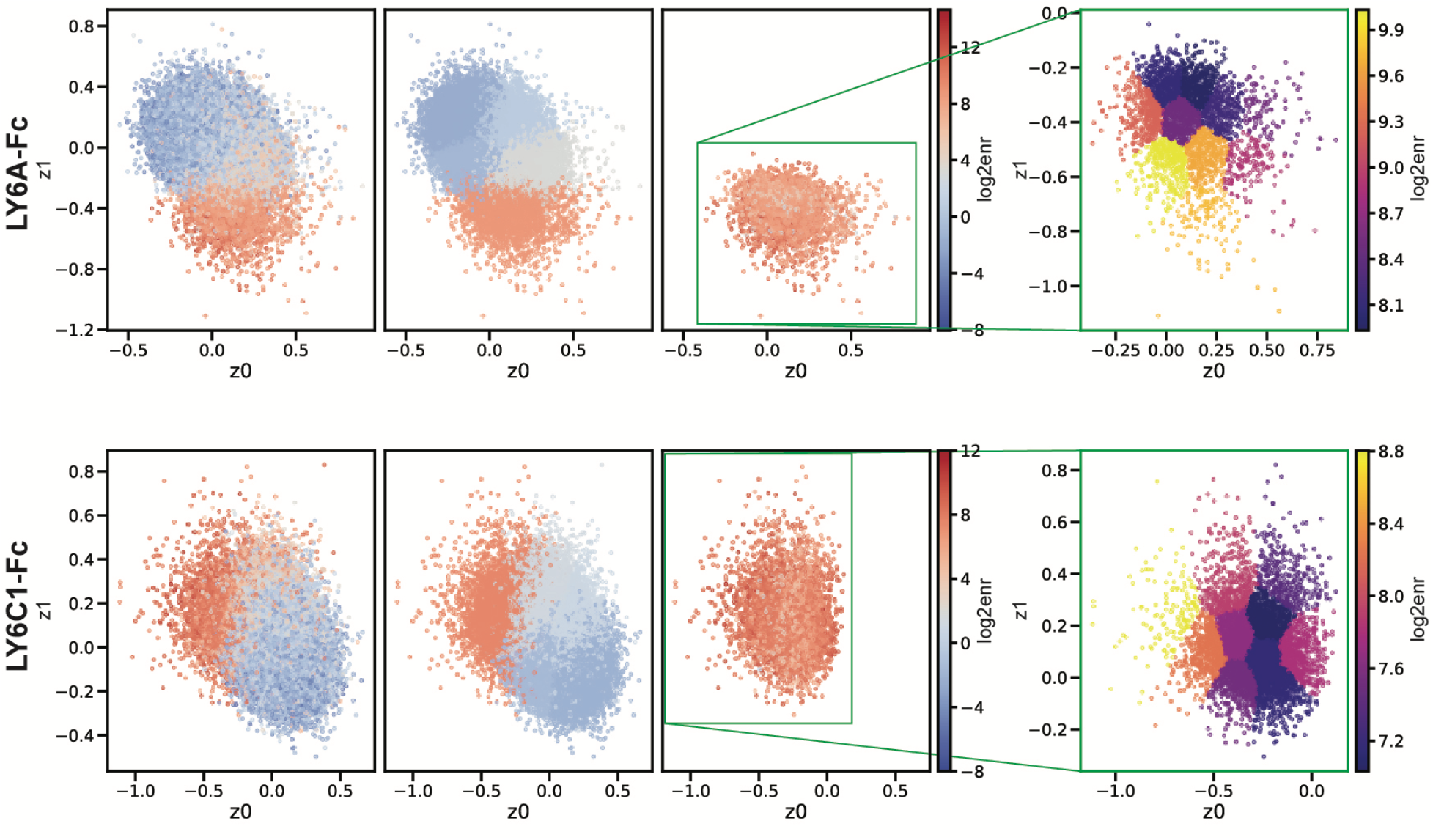
Plots of the LY6A-Fc and LY6C1-Fc training points in latent space. From left to right: all training points colored by assay log_2_ enrichment; all training points colored by mean primary cluster (see Methods, SVAE Sequence Generation) log_2_ enrichment; top (highest mean enrichment) cluster colored by assay log_2_ enrichment; top cluster further clustered into subclusters, colored by mean subcluster log_2_ enrichment. The first three plots from the left share spatial axes and color scale; the rightmost subclustering plot is centered on its own axes and recolored on its own scale.

**Figure 4—figure supplement 6.**
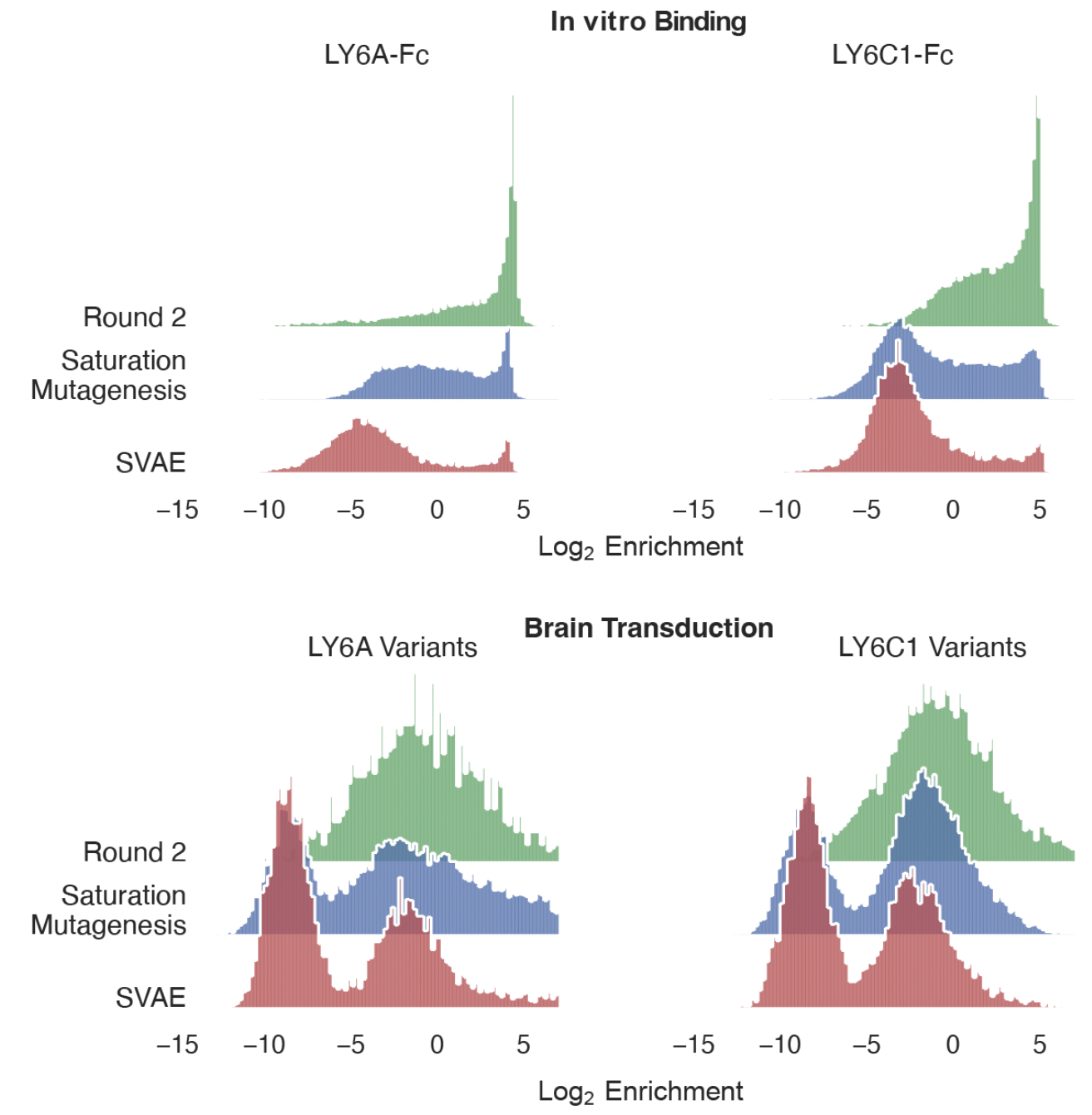
The *in vitro* binding and brain transduction enrichment scores of individual (nonclustered) Round 2, saturation mutagenesis, and SVAE variants. The data from Figure 4F is shown without clustering. The panels show the performance of variants for each assay: LY6A-Fc pull-down, LY6C1-Fc pull-down, C57BL/6J mouse brain transduction by LY6A-binding variants, or C57BL/6J mouse brain transduction by LY6C1-binding variants. The log_2_ enrichment of variants in each assay are shown and normalized so that the bar heights sum to 1 (proportion of variants).

